# Compensatory Plasticity Defines a Vulnerable Neuronal State in Alzheimer’s Disease with Psychosis

**DOI:** 10.1101/2025.04.30.651435

**Authors:** Matheus B. Victor, Na Sun, Kyriakitsa Galani, Noelle Leary, Yosuke Tanigawa, Aine Ni Scannail, Li-Lun Ho, Shaniah Prosper, Liwang Liu, Julia K. Kofler, Robert A. Sweet, Li-Huei Tsai, Manolis Kellis

**Affiliations:** Picower Institute for Learning and Memory, Massachusetts Institute of Technology, Cambridge, MA, USA; Department of Brain and Cognitive Sciences, Massachusetts Institute of Technology, Cambridge, MA, USA; MIT Computer Science and Artificial Intelligence Laboratory, Cambridge, MA, USA; Broad Institute of MIT and Harvard, Cambridge, MA, USA; Department of Pathology, University of Pittsburgh, Pittsburgh, PA, USA; Center for Neuroscience and Department of Psychiatry University of Pittsburgh, Pittsburgh, PA, USA; Department of Neuroscience, Icahn School of Medicine at Mount Sinai, New York, NY, USA; Whitehead Institute for Biomedical Research, Cambridge, MA, USA

## Abstract

Approximately 40 percent of Alzheimer’s disease patients develop psychosis (AD+P), yet the molecular and cellular basis of these symptoms remains poorly understood. Here we profiled single-nucleus transcriptomes and epigenomes from 48 postmortem Alzheimer’s brains stratified by psychiatric diagnosis. Across cell types, AD+P was distinguished by transcriptional programs in upper-layer cortical pyramidal neurons consistent with re-engagement of developmental and structural plasticity pathways. These neurons exhibited greater loss in AD+P cortex, indicating that such programs emerge in a context of heightened vulnerability. Integrating these findings with functional perturbation screens in stem-cell-derived brain organoids, we found that activation of these programs alters cortico-cortical network connectivity and can exacerbate network dysfunction. Our data suggest that compensatory neuronal plasticity, shaped by glial inflammatory responses, may paradoxically contribute to circuit instability and selective vulnerability underlying neuropsychiatric symptoms in dementia.

**Highlights:**

- Cell-type- and brain-region-specific transcriptional changes in AD with psychosis (AD+P)
- Upper-layer pyramidal dysfunction and metabolic vulnerability marks the pathophysiology of AD+P
- Circuit wiring programs are evoked in AD+P as maladaptive compensatory responses
- AD+P-associated IL-6 signaling impairs neuronal network function in brain organoids

## Introduction

Alzheimer’s disease (AD) is characterized by progressive memory loss, language deficits, impaired cognitive abilities, and behavioral disturbances, all of which severely compromise an individual’s ability to perform daily activities^1–3^. Hallmark pathological features of AD include the accumulation of amyloid-beta plaques and tau neurofibrillary tangles, accompanied by glial inflammation and reductions in brain volume, particularly within the hippocampus and prefrontal cortex, regions integral to memory formation and executive functions^4,5^. These molecular and structural alterations are often preceded by changes in brain activity patterns, detectable through neuroimaging techniques, which manifest within regions harboring early pathological accumulations long before clinical symptoms arise^6^.

Therapeutic efforts in AD have predominantly focused on mitigating cognitive decline. However, current pharmacological treatments offer limited efficacy in halting or reversing cognitive impairment^7^. Given the complex and multifactorial nature of AD, it is essential to consider alternative therapeutic targets beyond cognition, particularly those that address non-cognitive symptoms of AD like psychosis, which includes hallucinations and delusions^8^. Mitigating these symptoms may significantly enhance the quality of life for patients and alleviate the burdens faced by caregivers.

Alzheimer’s disease with psychosis (AD+P) is associated with accelerated cognitive deterioration, a heightened burden of tau pathology, pronounced neurodegeneration, and increased mortality risk^9^. Alarmingly, current antipsychotic treatments are accompanied by a black box warning due to a documented rise in fatal outcomes, underscoring the urgent need for novel and effective therapeutic strategies^10,11^. Despite the high prevalence and severe impact of psychosis in AD^12^, the underlying neurobiological mechanisms remain poorly understood. Genetic studies of AD+P suggest a common neurobiological pathway rather than nonspecific consequences of widespread neurodegeneration^13–16^. Moreover, emerging evidence implicates the glutamatergic system in the development of a number of psychotic disorders, particularly through neuronal hyperactivity and resultant disruptions in functional connectivity^17,18^. Neuropsychiatric symptoms in AD, including psychosis, have been linked to altered neural circuitry and synaptic dysfunction^19,20^, potentially stemming from maladaptive compensatory mechanisms in response to progressive neurodegeneration^21,22^.

In this study, we utilized single-nucleus RNA sequencing (snRNA-seq) and single-nucleus assay for transposase-accessible chromatin sequencing (snATAC-seq) to investigate the cellular and molecular landscape associated with psychosis in AD. Our analyses reveal that surviving excitatory pyramidal neurons in the upper cortical layers of AD+P brains re-engage developmental genetic programs aimed at circuit assembly, perhaps as an attempt to remodel network connectivity amid ongoing neurodegeneration. Surprisingly, data from stem-cell-based models indicate that these compensatory responses may inadvertently exacerbate neurodegenerative pathology and network dysfunction, offering new insights into the paradoxical nature of neuronal plasticity in AD+P.

## Results

### 1. snRNA-seq atlas of PFC and hippocampus in Alzheimer’s disease with and without psychosis

To investigate molecular and cellular differences in AD individuals with and without psychosis (AD+P vs. AD), we performed single-nucleus RNA-seq (snRNA-seq) on the prefrontal cortex (PFC) and anterior hippocampus (AH) from 24 AD+P and 24 AD individuals (**Methods and Patient Manifest in Supplementary Table S1**). After quality control, we obtained 160,875 transcriptomes, identifying major cell types, including excitatory neurons, inhibitory neurons, astrocytes, oligodendrocytes, oligodendrocyte progenitor cells (OPCs), microglia, vascular cells, and ependymal cells (**Fig. 1A**, **Methods**). Cell types were annotated based on canonical marker gene expression (**Fig. 1B**), and we further classified subtypes within excitatory and inhibitory neurons (**Fig. 1A**). Excitatory neuron subtypes were transcriptionally distinct by cortical layers and brain regions, while inhibitory neuron subtypes were distinguishable based on marker gene expression (**Fig. 1A**).

**Figure 1.**
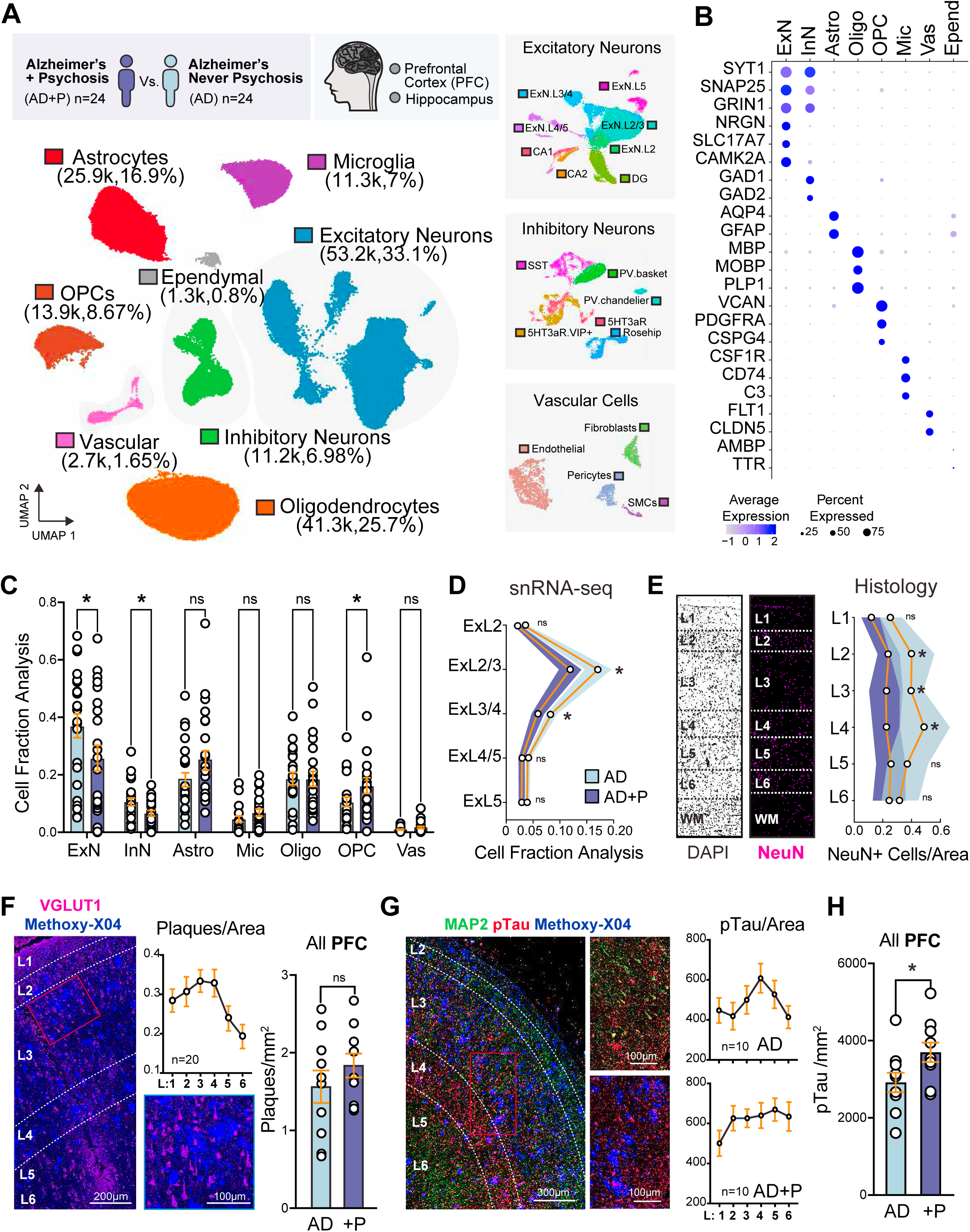
snRNA-seq atlas of PFC and hippocampus from AD patients with psychosis. **A.** Experimental design and UMAPs of 160,875 nuclei with annotated cell types in snRNA-seq data. The three UMAPs on the left show the high-resolution of cell types in excitatory neurons, inhibitory neurons and vasculature. **B.** Dotplot shows the expression of well-known marker genes for major cell types in the human brain. The color represents the expression level and the size of the dot represents the percentage of cells with expression. **C.** Cell fraction distribution of major cell types in AD w/wo psychosis. Propeller analysis to determine the statistical significance highlighted with *. **D.** Cell fraction distribution of excitatory neuron subtypes in AD w/wo psychosis. Propeller analysis to determine the statistical significance highlighted with *, (n= 24 AD-P and 24 AD+P; see Table S1 for patient manifest). **E.** Immunostaining for the neuronal marker NeuN, stratified by cortical layer based on nuclear density of DAPI (n= 10 AD-P and 10 AD+P). Two-Way ANOVA with Bonferroni post-hoc test. ns= not significant, *p-value <0.05. **F.** Immunostaining for the excitatory neuronal marker VGLUT1 as counterstain for the amyloid plaque stain, Methoxy-X04. Plaque density mapped across cortical layers (n=20 AD-P and AD+P combined). No overall significant difference in PFC, but near-significant trend for increased plaques in layer L4 based on Two-Way ANOVA with Bonferroni post-hoc test; plotted separately for panel only. **G.** Immunostaining for MAP2, Methoxy-X04 and phospho-tau (pTau, AT8). pTau density across cortical layers for AD-P and AD+P. **H.** Overall increase in pTau density in PFC (n= 10 AD-P and 10 AD+P); Student’s t-test, *p-value <0.05. Error bars represent S.E.M.

We then assessed whether specific cell type proportions differed between AD+P and AD individuals, accounting for confounding factors (**Methods**). In the PFC, we found that excitatory and inhibitory neurons showed significantly reduced proportions in AD+P, while OPCs were significantly increased in cortex (**Fig. 1C**). At the subtype level, we observed that cortical layer L2 and L3 (ExL2/3) and layers L3 and L4 (ExL3/4) excitatory neurons were disproportionately reduced in AD+P, suggesting cortical layer-specific neuronal loss associated with psychosis in AD (**Fig. 1D**). These correlates are based on marker gene expression that span layers 2 to 4 (L2-L4), suggesting that upper-layer excitatory pyramidal neurons may show differential vulnerability in AD+P. To confirm these findings based on high-throughput sequencing, we performed histology with immunohistochemistry (IHC) in a separate cohort consisting of tissue from 20 postmortem human brain PFCs (n= 10 AD+P versus 10 AD) (**Fig. 1E-H**, **Supplementary Table 2**). Images were segmented into layers based on nuclei density with the marker DAPI. Our histology analysis revealed consistent findings to our snRNA-seq cell fraction analysis, with AD+P subjects showing decreased density of neurons in upper cortical layers L2, L3 and L4, based on the expression of the pan-neuronal marker, NeuN (**Fig. 1E**). Neurons in deeper layers (i.e. L5 and L6) were not significantly altered between AD+P and AD. To understand how the distribution of pathology across the cortex correlates with neuronal loss, we quantified amyloid-beta plaques and hyperphosphorylated neurofibrillary tau tangles (NFTs) across PFC brain tissue (**Fig. 1F-H**). Numerous postmortem studies have reported that AD+P subjects exhibit greater NFT burden, while less consistent changes have been described for the levels of amyloid plaques^23–25^. We found that the density of amyloid plaques with the amyloid probe Methoxy-X04 are greater in upper cortical layers (L1-L4) and reduced in deeper layers (L5 and L6) across all subjects, suggesting upper cortical neurons bear greater plaque burden (**Fig. 1F**). Nevertheless, comparisons of amyloid densities segmented by psychosis presence (i.e. AD+P versus AD) did not show significant changes. When segmented by cortical layers, L4 showed a trend for increased plaques in AD+P (p-value = 0.09 with two-way ANOVA with bonferroni post hoc test) (**Fig. 1F and Supplemental Fig. S1A**). Next, we assessed if the levels of Tau phosphorylation (pTau), which is the main substrate of NFTs, were significantly altered in postmortem PFC brain tissue of AD+P relative to AD samples (**Fig. 1F**). Although the pattern of the distribution of pTau across the cortex were seemingly distinct between the two groups, we did not find statistically significant changes in specific cortical layers, but an overall increase across all layers (**Fig. 1F and Supplemental Fig. S1A**). These results suggest that neuropathological changes do not drive the differential vulnerability of upper cortical neurons in AD+P. However, our results do not rule out the possibility that neuropathological changes may act as a trigger to neuronal susceptibility. Of note, we did not find significant differences in the cell proportions of the AH (**Supplemental Fig. S2**). Although the mechanism at play remains unknown, we reasoned that transcriptional programs may reveal the differential responses in AD+P that lead to the relative greater loss of upper cortical neurons.

### 2. Cell-type- and brain-region-specific transcriptional changes in AD with psychosis

We next examined transcriptional changes in AD+P versus AD individuals by performing differential gene expression analysis for each cell type and brain region. Using MAST (single-cell level) and EdgeR (pseudo-bulk level), we obtained consistent results across methods (**Supplemental Fig. S3A-B, Methods**). In PFC, we identified 2,013 upregulated and 1,743 downregulated genes in AD+P individuals, with high cell-type specificity (**Fig. 2A**, **Supplemental Table 3**). In the hippocampus (AH), we found 932 upregulated and 1,012 downregulated genes (**Fig. 2B**, **Supplemental Table 4**), fewer than in the PFC, suggesting more dramatic transcriptional changes in the PFC at the time of sample collection.

**Figure 2.**
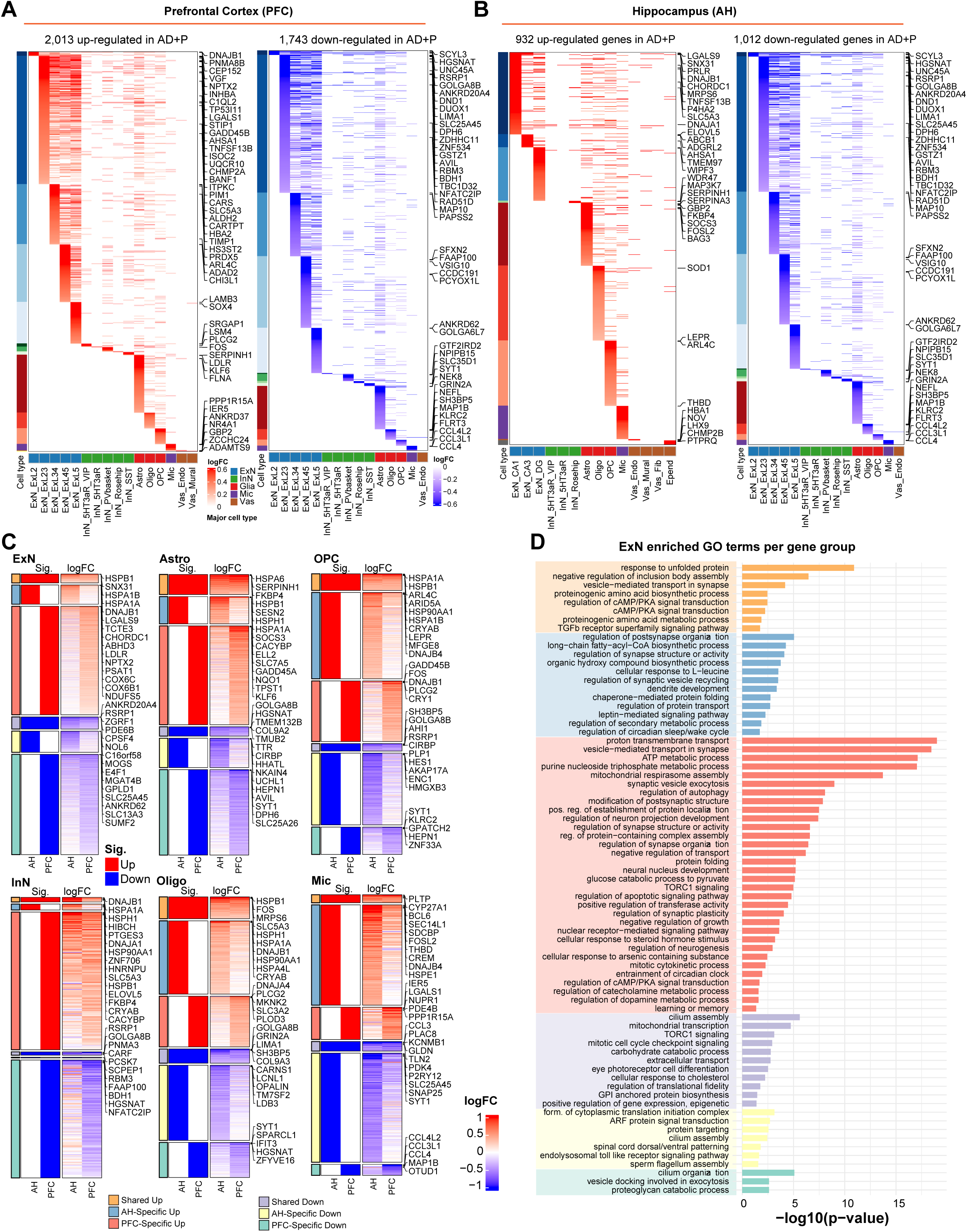
Cell-type-specific and brain-region-specific transcriptional changes in AD with psychosis. **A.** Heamaps show the log fold change of up-regulated and down-regulated genes per cell type in prefrontal cortex (SF). The top 5 DEGs per cell type are highlighted on the right side of the heatmap. **B.** Heamaps show the log fold change of up-regulated and down-regulated genes per cell type in hippocampus (AH). The top 5 DEGs per cell type are highlighted on the right side of the heatmap. **C.** The comparison of DEGs in AD+P between AH and SF per major cell type. Six groups of genes represent up-regulated and down-regulation sharing by two regions or unique to one region. For each heatmap, the left half shows whether the DEGs are significant (1 or 0) and the right half shows the real log fold change to present whether the change suggests the same direction between two regions. **D.** Significantly enriched and representative Gene Ontology (GO) terms in excitatory neurons binned in six groups based on shared or unique expression across brain tissue.

We systematically compared DEGs between the two regions and categorized them into six groups: shared upregulated, AH-specific upregulated, PFC-specific upregulated, shared downregulated, AH-specific downregulated, and PFC-specific downregulated. Gene expression changes showed consistent trends across regions, even with varying significance levels (**Fig. 2C**). Notably, the PFC had more region-specific DEGs in excitatory neurons, inhibitory neurons, and astrocytes, while the hippocampus showed higher microglia-specific DEGs. These findings suggest more severe neuronal loss in the PFC, consistent with the cell proportion analysis (**Fig. 1C**), and stronger microglia-driven inflammation in the hippocampus in AD+P individuals.

To explore the biological functions of DEGs in these six groups, we performed gene ontology (GO) enrichment analysis for each cell type (**Fig. 2C**, **Supplemental Fig. S4, Methods**). In excitatory neurons (**Fig. 2D**), we found that shared upregulated genes were enriched in protein unfolding and vesicle mediated transport in synapse. Hippocampus-specific upregulated genes were linked to protein folding and regulation of postsynapse organization, while PFC-specific upregulated genes were associated with synapse organization, neuron projection regulation, and regulation of apoptotic signaling. Shared downregulated genes were involved in cell cycle signaling and extracellular transport, while PFC-specific downregulated genes showed enrichment in cilium organization and signal transduction. These results highlight distinct and shared functional pathways in the PFC and hippocampus, emphasizing region-specific mechanisms in AD+P individuals.

Our cell fraction analysis and transcriptional results likely reflect the regional impact of AD progression, with more severe neurodegenerative phenotypes in AH than PFC^26^. Notably, our snRNA-seq profiling data was derived from subjects with an average age at death of 86 years old (ranging from 65 to 102 years old and balanced between AD+P and AD). Adding further support to the notion that disease progression may hinder our ability to uncover early pathological triggers, particularly as it relates to AH. It is also possible that the neural substrates that drive psychosis are indeed more dependent on cortical than hippocampal processes, particularly as it pertains to AD patients with or without psychosis. Nevertheless, due to the salience of our results in the PFC, we decided to focus on these processes to further derive mechanistic insights into the neural substrates associated with psychotic symptoms in dementia.

### 3. Compensatory mechanism evoked by surviving excitatory neurons in PFC

Neuronal DEGs in the PFC are particularly striking for pathways related to metabolic processes. This includes GO terms associated with up-regulated genes that relate to glucose and ATP metabolism, and mitochondrial respiration (**Fig. 2D**). Along with these transcriptional programs that may suggest metabolic dysfunction, excitatory neurons also exhibit up-regulated terms related to synaptic structure, organization and plasticity. To define the relative enrichment of these synaptic-related DEGs, we curated the genes from these GO terms and mapped their expression across cortical layers. Here, we stratified their relative enrichment by cortical laminae based on the co-expression of layer-specific markers. We found that synaptic-structure related DEGs were mostly enriched in upper cortical layers L2 and L3 (**Table S3**). This initial finding was counterintuitive, given that based on cell fraction analysis and histology we identified a significantly lower number of neurons in cortical layers L2 and L3 of the AD+P brains.

We hypothesized that the upregulation of genes related to synaptic structure and neuronal projection may rather reflect a compensatory mechanism by surviving neurons in the face of neurodegeneration. To test this idea, we probed PFC human brain sections from AD+P and AD-P subjects with fluorescent in situ RNA hybridization using an RNAscope probe for synapsin 1 (SYN1), an AD+P up-regulated DEG enriched in L2 and L3 which encodes a critical neuronal protein well-known to regulate synaptogenesis^27^ (**Supplemental Fig. 5A-B**). Notably, SYN1 is robustly and restrictively expressed by neurons^28^. Our analysis revealed that the number of RNAscope dots per nucleus for a given SYN1-positive neuron (comprising at least 3 SYN1 dots) increased in AD+P (**Supplemental Fig. S5C and S5D**). While these results were only significantly increased in L2 and L3, deeper layers also showed trends for increased SYN1 RNAscope dots. In agreement with our previous observations, the overall percentage of cells positive for SYN1 in cortical layers L2 and L3 were significantly reduced (**Supplemental Fig. S5E**). From these experiments we concluded that the relatively greater expression of this synaptic-related transcriptional signature in AD+P PFC captured by our snRNA-profiling is indicative of a compensatory mechanism by surviving neurons, perhaps in an attempt to maintain circuit function and integrity. Nevertheless, it remains unclear from these observations what are the impact of these genetic programs on the function of neural circuits.

The balance between excitation and inhibition (E/I) is tightly regulated in the adult cortex^29^. Disturbances to E/I balance are linked to many neuropsychiatric disorders, including schizophrenia ^30^. The association between schizophrenia and E/I disturbances has been implicated particularly with changes in the activity of Parvalbumin (PV) interneurons^31–33^. PV interneuron’s fast-spiking properties and fast GABA release are thought to be critical in shaping the activity of cortical ensembles and network synchrony^34^. PV-expressing interneurons can be divided into several subpopulations, with the most prominent being basket cells^35^. In our snRNA-seq analysis, PV basket cells are significantly decreased in AD+P, with DEGs indicative of widespread metabolic dysfunction (**Supplemental Fig. S5F and S5G**). Lack of balance in the ratio of excitatory to inhibitory neurons or perturbations in the activity of inhibitory neurons can greatly modify the output of excitatory neurons^36^. Many studies have also reported through functional imaging that AD patients with psychotic symptoms have reduced correlates of neuronal activity^8^. Hence, the functional consequence of our snRNA-seq results to circuit function is uncertain. Next, we decided to explore the functional impact of the gene expression perturbations we uncovered in AD+P brains to the activity of neuronal ensembles by employing induced pluripotent stem cell (iPSC)-based models of the human brain.

### 4. Functional characterization of AD with psychosis DEGs from upper-layer pyramidal neurons

The mammalian neocortex processes signals through local microcircuits that are arranged in distinct layers of information processing pathways^37^. Excitatory pyramidal neurons in layers L2 and L3 send long-range axons to distant brain regions, making them essential for cortical information processing^38^. We envisioned building an in vitro platform of cortico-cortical connectivity from which we could infer how genetic perturbations in L2 and L3 of AD+P brains impact circuit function. Human iPSCs patterned to adopt neural fates and forced to grow in 3D form structures known as brain organoids that have become sophisticated tools to model the brain in a human-based context^38,39^. Here, we employed established protocols to generate forebrain organoids^40^, composed predominantly of excitatory neurons, that recapitulate major cortical layers to engineer an input-output model of neuronal connectivity.

To gain control over neuronal activity patterns, we first generated an iPSC line that constitutively expressed the channelrhodopsin variant CheRiff ^41^ under the neuronal-restrictive promoter human Synapsin (hSyn-CheRiff). We then differentiated this line into forebrain organoids (**Fig. 3A**) We found that CheRiff and its increased sensitivity to blue light, allowed for faithful control of neuronal activity in organoids with blue light stimulation despite the high refractive index of culture media (**Fig. 3B**). Towards our goal of building a model of cortico-cortical connectivity, we created an assembloid by fusing two forebrain organoids together. Assembloids have been previously used to model connectivity across distinct brain regions, as they allow for organoids derived from multiple iPSC lines or organoids separately patterned to adopt distinct brain regionalities to be assembled post-production ^40^. To test the extent to which excitatory neurons in one organoid hemisphere projects across to the contralateral hemisphere, we infected organoids with the axonal-restricted genetically-encoded calcium indicator AAV Axonal-GCaMP6s and later generated assembloids with non-infected organoids^42^. This allowed us to image the activity of neurons whose projections crossed the assembloid junction line and confirm their input to the contralateral side (**Fig. 3C** and **Supplemental Movie S1**).

**Figure 3.**
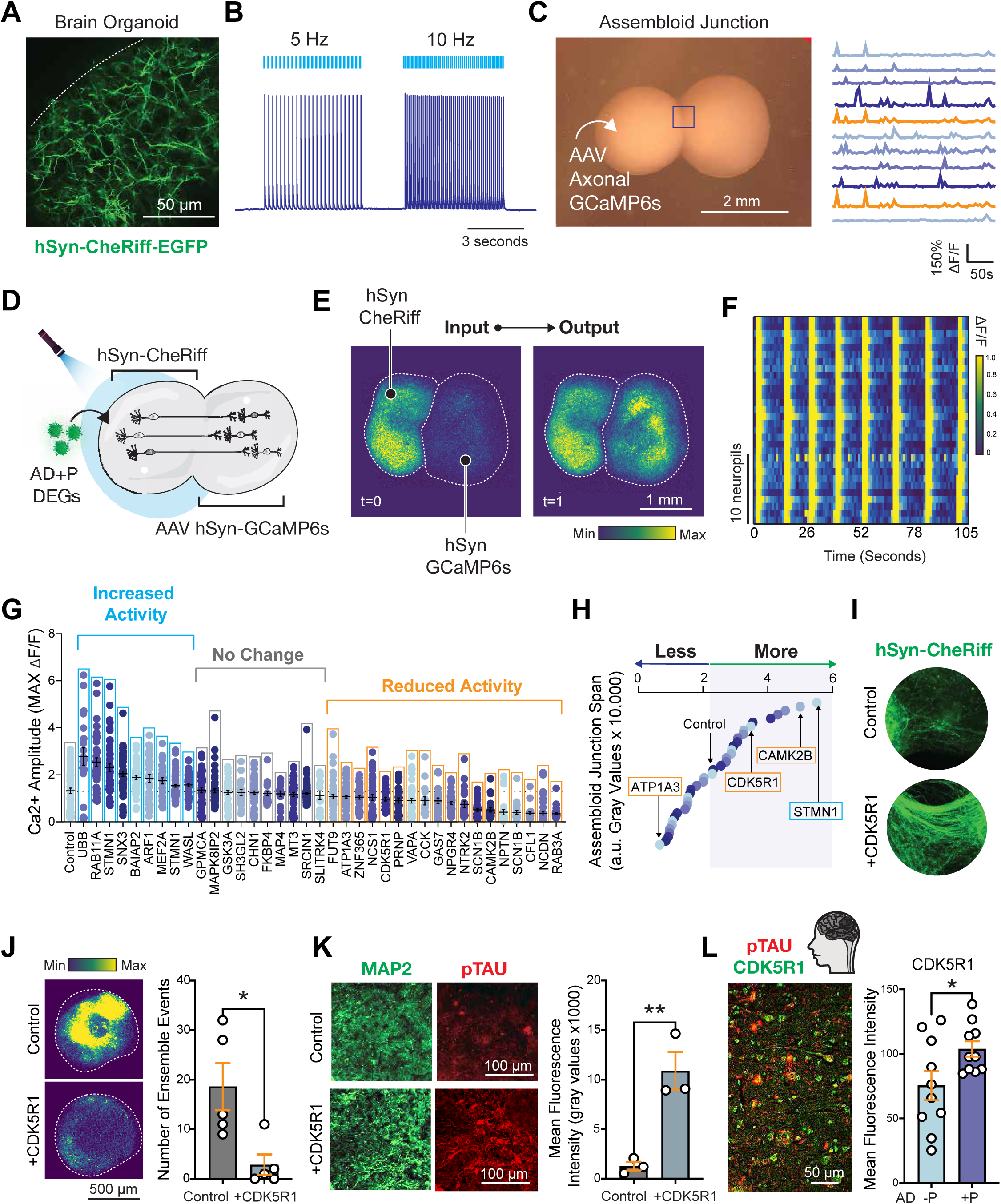
Functional Characterization of AD+P DEGs from Upper-Layers Excitatory Neurons in iPSC-derived Brain Organoids. **A.** Outer surface of an organoid derived from an iPSC line that constitutively expresses the channelrhodopsin variant CheRiff under the neuronal-restrictive promoter human Synapsin (hSyn-CheRiff). **B.** Whole-cell patch-clamp electrophysiology of iPSC-derived neurons harboring hSyn-CheRiff paced at 5Hz and 10Hz with blue light stimulation. **C.** AAV Axonal GCaMP6s infected organoids were fused with non-infected organoids to generate assembloids. Neuronal activity was assessed via calcium-imaging at the assembloid junction line, and calcium traces are shown for axonal spikes as the change in fluorescence over baseline fluorescence (ΔF/F). **D.** Assembloid input-output connectivity model with optogenetic stimulation to drive neuronal activity in one hemisphere and calcium imaging to readout for neuronal activity in the other hemisphere. **E.** Live-imaging of assembloids showing high-amplitude calcium events in the contralateral organoid. Color gradient represents minimum (purple) and maximum (yellow) levels of calcium amplitude. **F.** Rasterplot showing highly coordinated ensemble calcium activity dynamics. Change in fluorescence shown as Delta F/F. **G.** Assembloid quantification of contralateral calcium dynamics in response to optogenetic stimulation to the ipsilateral organoid overexpressing 37 genes from up-regulated DEGs in AD+P pyramidal upper-cortical layer neurons (L2 and L3) from snRNA-seq data. Data was ranked and binned into 3 groups (increased activity, no change, or reduced activity) based on Two-Way ANOVA with Bonferroni post-hoc test results. Calcium traces from the time series were manually segmented on ImageJ into individual cells based on threshold intensity results in n=15-88 neuropils per organoid and plotted for maximum amplitude. Control assembloids were transduced with a lentiviral vector encoding the blue fluorescent protein EBFP. **H**. Quantification of EGFP intensity on the contralateral organoid that traversed the assembloid junction line from hSyn-CheRiff ipsilateral organoid ranked from least to most fluorescent intensity. **I**. Representative image of hSyn-CheRiff across junction line from control and CDK5R1 overexpressing assembloid. **J.** Calcium dynamics in organoids 14 days post infection with CDK5R1 or EBFP lentivirus. Student’s t-test, *p-value <0.05, n= 5 organoids per group. **K**. Quantification of phospho-Tau (pTau) fluorescence levels in organoids infected with CDK5R1 or EBFP controls. Student’s t-test, **p-value <0.01, n= 3 organoids per group. **L**. Immunohistochemistry in postmortem brains showing increased CDK5R1 fluorescent intensity in PFC (n= 10 AD-P and 10 AD+P); Student’s t-test, *p-value <0.05. Error bars represent S.E.M.

Next, we fused organoids that harbored constitutive Syn-CheRiff expression with organoids previously infected with Synapsin-GCaMP6s to generate assembloids. Here, we would gain optogenetic control of neuronal activity in one hemisphere while having the capability of observing neuronal activity in the contralateral hemisphere via calcium imaging. Using this simple input-output model of neuronal connectivity, we reasoned that we could ectopically express synaptic-related upregulated DEGs from L2 and L3 of AD+P brains to the input hemisphere to probe its impact on the function and connectivity of neural circuits (**Fig. 3D**). In our model, optogenetic stimulation produced large amplitude calcium events on the contralateral organoid (**Fig. 3E**, **Fig. 3F**, **and Supplemental Movie S2**). We curated 37 genes upregulated in AD+P L2/L3 excitatory neurons from GO terms related to neuron projection development (genelist in **Table. S3**) and delivered them with lentiviral vectors under a ubiquitous promoter to hSyn-CheRiff organoids at 90 days after differentiation. In parallel, a separate set of organoids from the same line and batch that did not carry the hSyn-CheRiff construct were infected with AAV Synapsin-GCaMP6s. On day 120, these two sets of organoids were fused and made into assembloids. Ten days later, we screened activity patterns following stimulation in distinct assembloids in comparison to non-transduced controls (**Supplemental Movie S3**). We ranked the activity of the contralateral organoid based on calcium dynamics and binned them into groups of increased activity (n=9 of DEGs tested), no change (n=10 of DEGs tested), or reduced activity (n=18 of DEGs tested) in comparison to controls (**Fig. 4G**).

**Figure 4.**
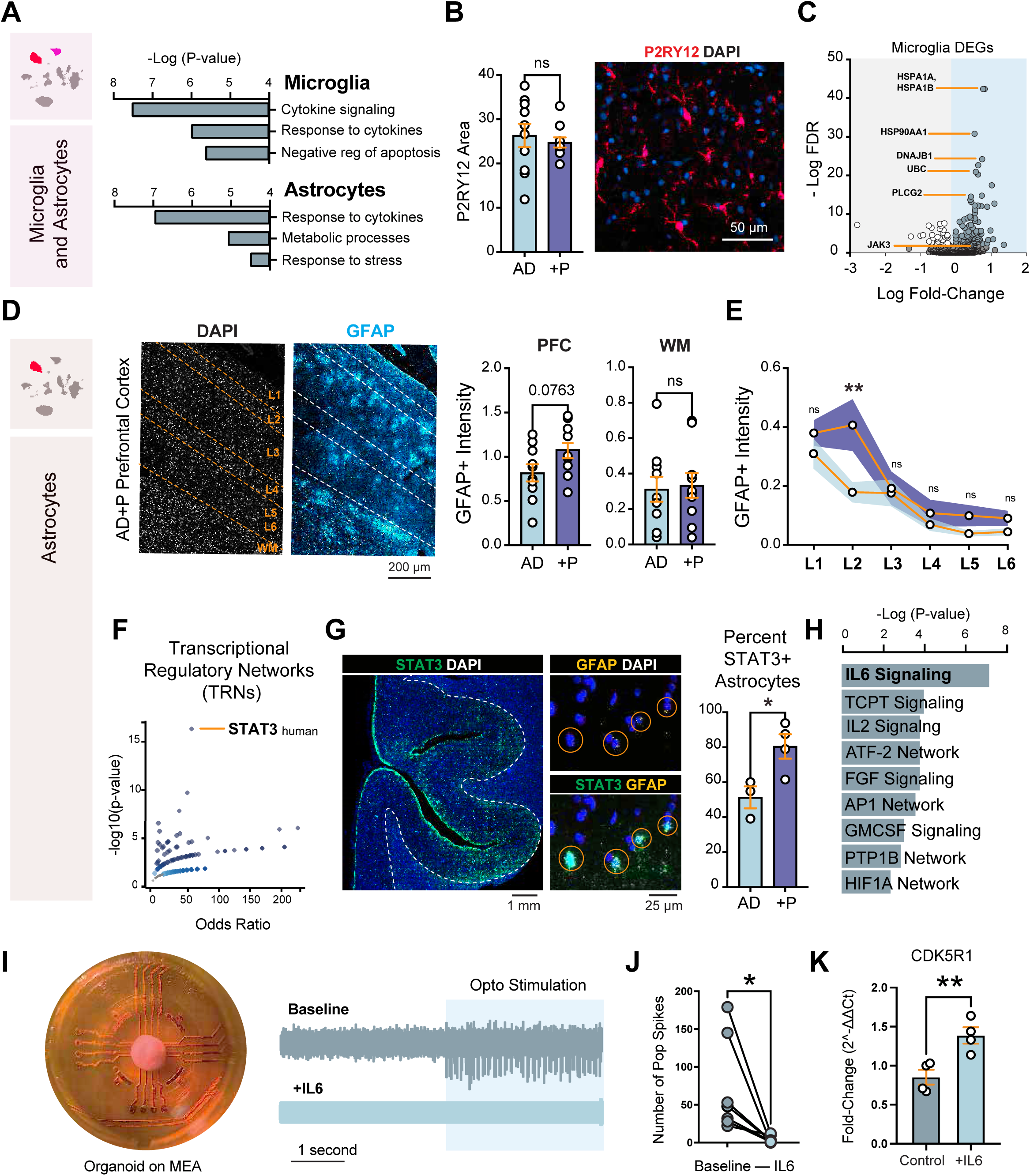
Glial Response in AD+P PFC is Marked by Microglial Exhaustion and Astrocyte-Driven Inflammation. **A.** Representative GO terms for microglia and astrocytes from PFC DEGs. **B.** Histology for microglia-marker P2RY12 in PFC. Student’s t-test, ns= not significant, n= 10 AD-P and 10 AD+P. **C.** Volcano plot for up and down-regulated genes in AD+P PFC microglia. **D.** Immunofluorescence with the atrocyte-marker GFAP quantified across PFC cortical layers and white matter (wm) normalized per area. Student’s t-test, ns= not significant, n= 10 AD-P and 10 AD+P postmortem brains. **E.** GFAP quantification binned by cortical layers. Two-Way ANOVA with Bonferroni post-hoc test. ns= not significant, **p-value <0.01, n= 10 AD-P and 10 AD+P postmortem brains. **F.** Transcriptional Regulatory Network (TRN) from astrocyte top GO term “cellular response to cytokine stimulus”. **G**. RNAscope for STAT3 and GFAP. (n= 3 AD-P and 4 AD+P); Student’s t-test, *p-value <0.05. H. Ranked Pathway Interaction Database (PID) Pathways. **I.** hSyn-CheRiff organoid seeded on multielectrode array (MEA) and raw traces from recordings of the same organoid at baseline and after 5 ng/ml of IL-6. **J.** Paired t-test, n= 7 organoids recorded at baseline and again following overnight incubation with IL-6, *p-value <0.05. **K.** qPCR for CDK5R1 following treatment of 5 ng/ml of IL-6. (n= 4 organoids per group); Student’s t-test, **p-value <0.01. Error bars represent S.E.M.

Given that we chose to ectopically express genes related to neuronal projection development, and that the majority of the DEGs we tested decreased activity on the contralateral organoid, we wondered to what extent these genetic manipulations modify the span of projections across assembloid junctions. Using the stable GFP signal of the hSyn-CheRiff line, we traced fluorescence to the contralateral hemisphere near the vicinity of the assembloid junction and ranked it from least to most projection (**Fig. 3H**). Interestingly, we found instances of DEGs that decreased activity on the contralateral organoid but counterintuitively increased projections across the assembloid junction. One such example is CAMK2B, a gene that encodes the beta subunit of the calcium/calmodulin-dependent protein kinase II and plays a critical role in synaptic plasticity and dendritic arborization^43^. In experiments overexpressing CAMK2B or an identical construct that instead carried the blue fluorescent protein EBFP, we found that CAMK2B overexpression leads to enhanced vulnerability of neurons following optogenetic stimulation compared to controls, likely due to excitotoxicity (**Supplemental Figure S6**). In fact, overexpression of CaMKII has previously been shown to increase glutamate-induced neuronal death in similar contexts^43,44^. While the overexpression of CDK5R1, an AD+P L2/L3 upregulated DEG, led to similar phenotypes as CAMK2B overexpression (i.e. less contralateral activity but more projection across assembloid junction), we did not find evidence of enhanced excitotoxicity (**Fig. 3H** and **Supplemental Figure S6**). CDK5R1 encodes for a protein known as p35, a neuron-specific activator of the cyclin-dependent kinase 5 (CDK5)^45^.

While the activation of CDK5 is required for proper central nervous system development and cortical lamination ^45,46^, CDK5 is also a key enzyme responsible for the phosphorylation of tau^47^. Of note, p35 can be proteolytically cleaved by excitotoxic triggers such as amyloid beta aggregation or high levels of intracellular calcium to generate a smaller peptide known as p25^48,49^. Interestingly, the p25 form of this protein has been shown to accumulate in neurons of patients with AD^50^. Since p25 has been shown to increase CDK5 kinase activity, it is thought that this could be a critical mechanism linking disease-associated toxic events to neuronal tau hyperphosphorylation and the pathological formation of NFTs^51^. In mice, CDK5R1 peaks in expression during early development where it plays a crucial role in promoting neurite outgrowth^51,52^, congruent with our observation of greater projections across the assembloid junctions in CDK5R1 overexpressing organoids. Since we observed decreased activity on the contralateral organoid, we decided to test the impact of CDK5R1 overexpression on the infected organoid itself. We found that CDK5R1 overexpression dramatically decreased levels of neuronal activity in infected organoids two weeks post-transduction (**Fig. 3I and 3J**).

To determine the integrity of the neurons following CDK5R1 overexpression, we fixed and performed immunostaining on infected organoids derived from a healthy control subject cultured for over 18 months. Staining for the neuronal marker MAP2 showed neuronal processes that were free of blebbing with intact morphologies (**Fig. 3K**). We probed pTau at a known Cdk5 phosphorylation site, Ser396 ^53^, in CDK5R1 overexpressing or EBFP-infected organoids and found a robust increase in the levels of pTau with CDK5R1 overexpression (**Fig. 3K**). It is important to also note that many of the Cdk5 target sites are hyperphosphorylated in post-mortem AD brain samples^51^. Since we had already established the greater pTau burden with IHC in a separate cohort of postmortem human brain PFCs (**Fig. 1F and Supplemental Fig. S1A)**, we next probed the same tissue for CDK5R1 levels. Concordant with the increase in CDK5R1 transcript levels in our snRNA-seq dataset, we also observed higher CDK5R1 protein expression in AD+P compared to AD subjects. (n= 10 AD+P versus 10 AD) (**Fig. 3L**). Taken together, our findings point to a compensatory shift in pyramidal neurons marked by enhanced structural plasticity and metabolic load, which could represent an effort to preserve circuit function in the face of neurodegeneration. Yet our in vitro experiments with repeated optogenetic stimulation suggest that this response may come at a cost, predisposing these neurons to greater vulnerability under stress. Notably, many functional imaging studies to date have reported that AD patients with neuropsychiatric symptoms have decreased gray matter volume and reduced network connectivity^54^. Nevertheless, the molecular and cellular cues that evokes these genetic programs in AD+P excitatory upper-layer neurons leading to their preferential vulnerability remains unclear.

### 5. Glial response in AD with psychosis is marked by microglial exhaustion and astrocyte-driven inflammation

Next we sought to explore how transcriptional programs in glia may reflect our findings of neuronal vulnerability in the AD+P cortex. To preserve sensitivity to state-dependent transcriptional changes within heterogeneous glial populations, we focused on single-cell level differential expression using MAST for all subsequent glial analyses. This approach allowed us to capture glial activation and response programs that would likely be diluted by pseudobulk averaging across mixed cellular states. Here, we were particularly interested in exploring the crosstalk of astrocytes and microglia, as the most significant pathways for DEGs in each cell type suggested enrichment of an inflammatory signaling axis. Up-regulated DEGs for astrocytes include GO terms for “cellular response to cytokine stimulus” while up-regulated DEGs for microglia include “cytokine-mediated signaling pathway” (**Fig. 4A** and **Supplemental Table S5**). Microglial transcriptional programs in response to aging and AD have been previously deeply characterized with snRNA-seq from mouse models of neurodegeneration and postmortem human brains^55–59^. While a wide variety of transcriptional programs associated with disease states have been catalogued, microglial activation across disease is consistently marked by the downregulation of a handful of homeostatic gene markers. One such case is the loss of P2RY12 expression, a purinergic receptor thought to mediate microglial surveillance^60^. Even though our cell fraction analysis did not capture any significant changes in microglial content between AD+P and AD brains (**Fig. 1A**), we attempted to determine via immunohistochemistry if the coverage of P2RY12-positive microglial processes were changed between AD+P and AD subjects. Our analysis did not find significant changes (**Fig. 4B**), suggesting that microglia number and activation status is similar across AD independent of neuropsychiatric diagnosis. At closer inspection, up-regulated DEGs of microglia revealed a widespread genetic program reflective of cellular response to stress (e.g. unfolded proteins and heat shock proteins) (**Fig. 4C** and **Supplemental Table S5**). Additionally, upregulated DEGs include PLCG2, whose expression increases in plaque-associated microglia^61^, suggesting that perhaps AD+P stress-induced microglial states could be triggered by neuropathology burden. Moreover, AD+P microglia differentially upregulated canonical markers of inflammation, such as JAK-STAT pathway (e.g. JAK3 and STAT3^62^). Interestingly, upregulation of stress and inflammatory programs in AD+P microglia is also accompanied by well-established markers of cellular senescence (i.e. CDKN1A which encodes the protein p21)^63^ (**Supplemental Table S5**). Many reports have now identified that exposure to tau is a potent inducer of microglial senescence^64,65^. Hence in the AD+P cortex, microglia adopt an exhausted state that is marked by pro-inflammatory programs, possibly reflecting a maladaptive response to persistent stress signals in the local microenvironment. This may represent a decoupling of classical neuropathology burden from immune cell activation, where microglia respond to subtle cues, such as altered synaptic function or tau pathology, rather than overt plaque deposition.

While our cell fraction analysis did not capture changes in astrocytic content between AD+P and AD brains, there was a non-significant trend for increased numbers of astrocytes in AD+P subjects (**Fig. 1A**), which was similarly the case through histology of cortical gray matter using the astrocyte marker GFAP (**Fig. 4D**) (n= 10 AD+P versus 10 AD; p-value = 0.076, student’s t-test). Using the nuclear stain DAPI to parse cortical layers, our quantification of astrocytes across the cortical gray matter revealed greater levels of GFAP specifically in layer L2 of AD+P brains (**Fig. 4E**). To further define the astrocytic response, we attempted to identify the master regulators of this transcriptional program and build a Transcriptional Regulatory Network (TRN)^66^. Here, we used the top GO term related to the up-regulated DEGs of “cellular response to cytokine stimulus” and found that STAT3 showed the greatest enrichment for transcription factors (TFs) (**Fig. 4F**). Of note, we found that STAT3 itself is upregulated in astrocytes of AD+P versus AD subjects (**Supplemental Table S5**). To confirm this finding, we probed PFC human brain sections from AD+P and AD subjects with fluorescent in situ RNA hybridization using RNAscope probes for STAT3 and GFAP (**Fig. 4G**). Using this approach, we first defined astrocytes as cells bearing at least 3 GFAP RNAscope dots, and subsequently mapped the percent of these cells that were also double positive for STAT3. We focused this analysis specifically on layers L2 and L3 since we had seen greater GFAP signal in the proximity of this cortical region in AD+P brains. Our results identified a robust increase in the presence of STAT3 expression in AD+P astrocytes, further confirming our snRNA-seq dataset (**Fig. 4G**).

To define the cellular signaling pathways that drive this response, we imputed the same list of up-regulated DEGs into the Pathway Interaction Database (PID)^66^ and found that IL6-mediated signaling is the major driver of this STAT3 astrocytic response (**Fig. 4H**). IL-6 (Interleukin-6) is a pleiotropic cytokine involved in the activation of the transcription factor STAT3 through JAK/STAT signaling^67^. Of interest, the IL6ST, a gene that encodes a critical interleukin 6 cytokine family signal transducer^67,68^, is also up-regulated in AD+P astrocytes (**Supplemental Table S5**). In the brain, IL-6 is largely produced by activated astrocytes and microglial cells^69^. Furthermore, aberrant IL-6 signaling is known to lead to neuronal dysfunction and synaptic loss and therefore to be correlated with cognitive decline and memory impairment in AD^70^. To test the impact of IL-6 signaling to neuronal physiology and CDK5R1 expression, we treated organoids seeded on multielectrode arrays (MEA) with recombinant IL-6 and recorded their activity (**Fig. 4I-4K**). Using organoids derived from our hSYN-CheRiff optogenetic iPSC line, we first tested the ability of optogenetic stimulation to drive population spikes following overnight incubation with 5ng/ml of recombinant IL-6. During a 15 minute recording session, we observed a dramatic decrease in activity in organoids with very few spikes detected upon 10Hz blue stimulation when treated with IL-6 (**Fig. 4I** and **Fig. 4J**). It has been previously reported that IL-6 treated primary mouse neurons accumulate abnormal levels of hyperphosphorylated tau, which is mediated through increased levels of CDK5R1 and its encoded protein p35, the activator of CDK5 ^71^. Hence, we tested the expression of CDK5R1 in IL-6 treated organoids and found that indeed suppression of neuronal activity is concomitant with up-regulation of CDK5R1 expression (**Fig. 4K**). Collectively, these results point to a neuroinflammatory axis driven by IL-6 that promotes astrocyte activation and the demise of circuit function.

### 6. Epigenetic regulation of cell-type-specific gene expression changes and association with Schizophrenia genetics

To uncover the epigenetic mechanisms underlying cell-type-specific gene expression changes, we analyzed chromatin accessibility using single-nucleus ATAC-seq (snATAC-seq) from human postmortem brain samples of the same cohort (**Methods**). From 241,430 high-quality nuclei, we identified epigenetic patterns across major brain cell types, including excitatory and inhibitory neurons, astrocytes, oligodendrocytes, OPCs, microglia, and vascular cells, using promoter signals of canonical marker genes (**Fig. 5A**, **Supplemental Fig. S7A**). Similar to snRNA-seq results, nuclei were clustered primarily by cell identity in UMAP space, regardless of phenotypic group or brain region (**Fig. 5B-C**). Cell proportion analysis revealed fewer neurons in the hippocampus compared to the PFC (**Supplemental Fig. S7B**). Notably, PFC snATAC-seq data captured fewer excitatory neuron nuclei than snRNA-seq data, possibly reflecting epigenetic erosion^72^ (**Supplemental Fig. S7C**).

**Figure 5.**
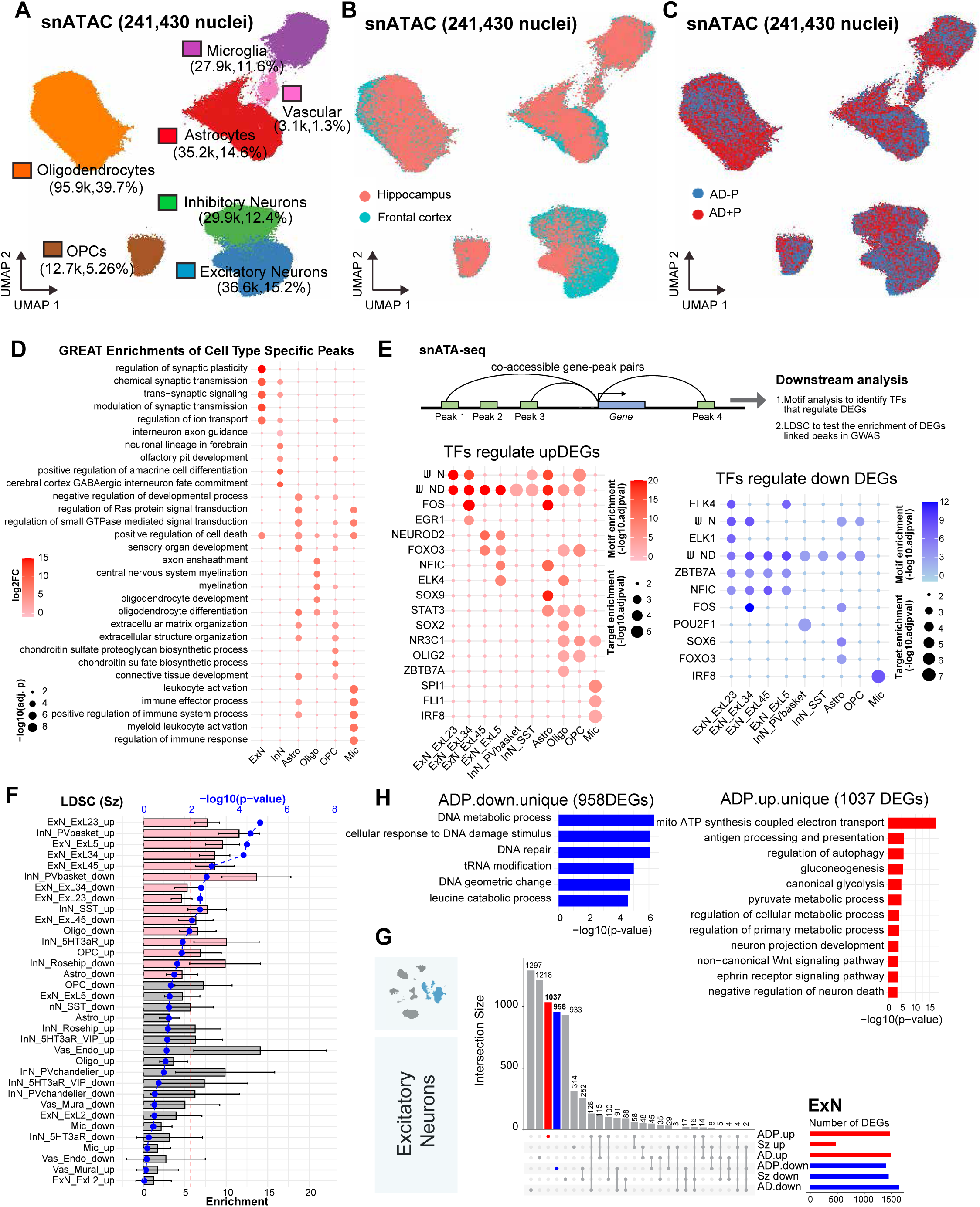
Epigenetic regulation of cell-type-specific gene expression changes and their association with genetics. **A-C.** UMAPs of 241,430 nuclei in snATAC-seq data with annotated cell types (**A**), brain regions (**B**) and pathological groups (**C**). **D.** GREAT enrichment analysis for cell type specific peaks to show the function relevance with certain cell types. The color represents the log2 fold change and the size of dots represents the significance. **E**. The diagram to show the co-accessible gene-peak pairs and their applications in TF identification and genetic association analysis by LDSC. TF enrichment for up-regulated DEGs and down-regulated DEGs per cell type. The dotplot shows the motif enrichment significance by color and TF-target enrichment significance by size of dots. **F.** AD+P DEGs linked chromatin accessible peaks show the heritability enrichment of genetic variants implicated for Schizophrenia using S-LDSC analysis. The barplot shows the enrichment score, where pink ones represent nominally significant cell types. The -log10 of p-value is shown by the blue point. The red dash line represents the stringent threshold with p-value < 0.01.**G**. Representative Gene Ontology terms enriched in AD+P unique down-regulated DEGs. **H**. Representative Gene Ontology terms enriched in AD+P unique up-regulated DEGs. **I**. The UpSet plot shows the overlap of DEGs between Alzheimer’s Disease (AD), Schizophrenia (Sz) and AD+P (ADP). The pink bars show the significant intersections. The blue and red bars show the AD+P specific down-/up-regulated DEGs. Error bars represent S.D.

To validate cell-type annotation in snATAC-seq, we integrated snRNA-seq and snATAC-seq data for each brain region, in which nuclei from the same annotated cell type clustered tightly together (**Supplemental Fig. S7D-E**). We identified accessible chromatin peaks, annotated their genomic features, and performed GREAT enrichment analysis, which revealed cell-type-specific biological functions (**Fig. 5D**, **Supplemental Fig. S7F, Supplementary Table S6**). For instance, peaks in excitatory neurons were enriched in synaptic plasticity, trans-synaptic signaling, and ion transport regulation. Inhibitory neuron peaks were enriched in interneuron axon guidance and fate commitment. Astrocyte peaks were linked to Ras signaling and sensory organ development. Oligodendrocyte peaks were associated with myelination and axon ensheathment, while microglial peaks were enriched in immune effector processes and regulation of immune response. These findings demonstrate that cell-type-specific epigenetic patterns reflect functional identities and provide insights into gene expression changes in disease contexts.

To explore how epigenetic regulation influences gene expression, we linked distal peaks to promoters using co-accessibility patterns (**Methods**). We identified 158,246 peak-to-promoter links implicating 16,193 genes and 82,695 peaks (**Supplemental Fig. S7G, Supplementary Table S7**). Motif analysis on distal peaks associated with differentially expressed genes between AD+P and AD-P individuals identified 171 transcription factors (TFs) regulating DEGs in SF (**Methods**). Additionally, we inferred TFs whose known targets were enriched among DEGs using enrichment analysis. By integrating these approaches with expression of TFs, we identified 17 TFs regulating upregulated DEGs in AD+P and 11 TFs regulating downregulated DEGs (**Fig. 5E**). Notably, TFs like FOS, JUND, and JUN were implicated in both upregulated and downregulated DEGs, indicating dual regulatory roles. Notably, SPI1 and IRF8, two key microglial transcription factors, were enriched among regulators of upregulated microglial genes, suggesting activation of immune and interferon-related pathways in AD+P. Interestingly, IRF8 also appeared among TFs linked to downregulated DEGs, indicating a dual regulatory role in both activating immune responses and repressing homeostatic gene networks. Together, these results highlight cell-type-specific and context-dependent TF activity shaping transcriptional reprogramming in Alzheimer’s disease with psychosis.

To contextualize the transcriptional differences distinguishing AD+P from AD, we directly compared our cell type specific DEGs with AD-associated signatures derived from the single-cell transcriptomic atlas of the aged human prefrontal cortex reported by Mathys et al^73^. To enable a direct and methodologically consistent comparison, we re-analyzed the Mathys dataset using the same differential expression framework applied in our study. This dataset, which profiles 2.3 million cells from 427 individuals spanning minimal to severe AD pathology, provides a high-resolution reference for canonical AD-related gene expression changes relative to individuals with limited pathology^73^. Across all major brain cell types, AD+P log fold changes showed little correlation with the direction or magnitude of AD-associated changes observed in the re-analyzed Mathys dataset, with most genes clustering near zero on both axes (**Supplemental Fig. S8**). The lack of a diagonal trend indicates that psychosis-related transcriptional changes do not simply reflect quantitative amplification of established AD expression programs. Consistent with this observation, Fisher’s exact tests revealed minimal overlap between AD+P DEGs and the cell type specific AD-upregulated or AD-downregulated gene sets from Mathys et al. (**Supplemental Fig. S9**). Together, these findings support a model in which psychosis in Alzheimer’s disease reflects cell-type-specific transcriptional programs superimposed on a broadly shared AD molecular landscape, rather than a uniform amplification of canonical AD pathways at the tissue level.

We further investigated how genetics influences downstream gene expression changes. Using Stratified Linkage Disequilibrium Score Regression (S-LDSC)^74^, we evaluated the heritability enrichment of AD and schizophrenia genetic variants in accessible peaks linked to DEGs in AD+P (**Methods**). Schizophrenia variants were more associated with AD+P DEGs than AD variants (**Fig. 5F**, **Supplemental Fig. S10A**), implying a shared genetic basis for psychotic symptoms in AD and schizophrenia. For instance, DEGs in layers 2/3, 3/4, and 4/5 excitatory neurons in AD+P were strongly associated with schizophrenia variants (**Fig. 5F**). Moreover, comparing DEGs across AD vs. controls, AD+P vs. AD, and schizophrenia vs. controls revealed shared downregulated DEGs in excitatory neurons between AD+P and schizophrenia (**Fig. 5G**, **Supplemental Fig. S10B-C**). Gene Ontology analysis showed AD+P-specific downregulated DEGs in excitatory neurons were enriched in DNA repair and DNA geometric change, while upregulated DEGs were linked to gluconeogenesis, neuron projection development, and regulation of autophagy (**Fig. 5H**). These findings highlight distinct molecular and cellular responses to psychosis in AD patients.

## Discussion

By integrating single-nucleus transcriptomic and epigenomic profiling of postmortem brains with functional studies in stem-cell-derived models, our work reveals that upper-layer cortical pyramidal neurons in AD+P engage transcriptional programs associated with structural remodeling and cellular stress. Although these signatures emerge in the setting of prominent glial inflammation and tau pathology, the temporal sequence of these events remains unresolved. Our data do not distinguish whether these neuronal transcriptional changes arise as downstream responses to pre-existing pathology or whether they contribute to circuit instability and selective vulnerability. Both AD and AD+P subjects exhibit dense amyloid and tau pathology in superficial cortical layers, providing a shared upstream context that may drive neuroinflammatory signaling and neuronal compensatory responses. In this framework, structural remodeling programs in excitatory neurons may represent an attempt to preserve connectivity under conditions of synaptic disruption. Conversely, such remodeling may occur in a context of heightened vulnerability, consistent with our observation of increased loss of these neurons in AD+P. Prior work has shown that upper-layer pyramidal neurons mount compensatory transcriptional responses early in AD^75^, but whether these programs reflect adaptive remodeling or early maladaptive states is not yet known.

Additionally, our data suggest that metabolic stress in interneurons may contribute to their vulnerability and impairments in inhibitory tone, which could influence circuit stability. Given that interneurons depend heavily on oxidative metabolism, disruptions in their energetic support, potentially mediated by glial inflammation or mitochondrial dysfunction, may compromise their ability to sustain inhibitory function^76^. Our findings also implicate microglial exhaustion and senescence in AD+P, which may further alter the cellular environment required for interneuron function. Microglia in AD+P exhibit loss of homeostatic gene expression and upregulation of stress and senescence markers such as CDKN1A (p21), suggesting diminished functional capacity despite preserved cell numbers. Prior studies have linked microglial loss-of-function to impaired regulation of neuronal networks^77–79^, raising the possibility that microglial senescence in AD+P contributes to an environment in which interneurons struggle to maintain inhibitory tone. This interplay between glial dysfunction and interneuron vulnerability may contribute to altered circuit dynamics in AD+P. Clarifying how these cell types interact under conditions of chronic inflammatory stress will be essential for identifying therapeutic strategies that restore network homeostasis in AD+P.

It is also important to note that hallucinations are a hallmark feature of Lewy body dementia (LBD), and their high prevalence often complicates the clinical distinction between LBD and AD+P. Although these are distinct disorders, mixed neuropathology is common among individuals with Alzheimer’s disease neuropathological changes (ADNC), with many patients exhibiting varying degrees of Lewy body pathology in addition to amyloid and tau lesions. This pattern is also reflected in our snRNAseq cohort: among patients clinically diagnosed with Alzheimer’s disease and confirmed as primary ADNC, neocortical Lewy body pathology was present in 9 of 24 AD+P patients. In contrast, our histology validation cohort was more balanced, with neocortical Lewy body pathology detected in only one AD+P case, though early-stage Lewy body involvement outside the neocortex was present in several individuals in both groups. Given the potential for alpha-synuclein, the defining proteinaceous component of Lewy bodies, to influence molecular signatures in our study, we included neocortical Lewy body status as a covariate in our differential expression analyses. Beyond controlling for this effect, the presence of mixed pathology in our cohort provides a valuable opportunity to explore transcriptional states associated with co-occurring Alzheimer’s and Lewy body pathology, offering broader insight into the molecular architecture of neurodegenerative comorbidity.

Recent bulk gray matter proteomics in dorsolateral prefrontal cortex reported that AD+P exhibits more differentially abundant proteins versus controls, but that AD+P and AD without psychosis show highly correlated case control proteomic changes, consistent with quantitative shifts within a shared AD proteomic landscape^80^. Our single-nucleus transcriptomic and epigenomic analyses refine this interpretation by resolving molecular changes at cellular resolution. While bulk tissue measurements capture dominant, shared disease processes, they can obscure state- and compartment-specific programs arising in discrete neuronal and glial populations. In this context, the AD+P-associated signatures we identified, particularly in upper-layer pyramidal neurons and glial response programs, likely represent transcriptional states that contribute to circuit vulnerability but are diluted in homogenate-level analyses. Notably, this recent proteomics study also identified synapse- and inflammation-associated protein modules correlated with postsynaptic density yield^80^, conceptually aligning with our cell-type-resolved evidence linking glial inflammatory responses to neuronal susceptibility. Together, these findings suggest that psychosis in AD emerges from selective cellular programs layered onto a common neurodegenerative framework.

Our functional studies suggest that the upregulation of CDK5R1, the regulatory subunit of CDK5, may be driven by glial inflammatory responses in AD+P. CDK5 is a key kinase involved in neuronal development and synaptic plasticity, but its dysregulation has also been implicated in tau hyperphosphorylation and neurodegeneration^51^. Therefore, CDK5R1 may act as a switch that couples attempted structural repair to biochemical changes that accelerate tau pathology and neuronal failure. Notably, inhibition of CDK5 has been shown to ameliorate tau pathology and prevent neuronal loss in preclinical models of AD^81^. Given the strong association between CDK5 activity, tau pathology, and circuit dysfunction in AD+P, our findings suggest that CDK5 inhibitors^82,83^ may hold promise for mitigating not only neurodegeneration but also the neuropsychiatric symptoms associated with AD. Future investigations should explore whether CDK5 inhibition can restore synaptic balance and attenuated psychotic symptoms in preclinical AD+P models, paving the way for targeted therapeutic interventions.

Additionally, we uncovered convergent and divergent molecular features of psychosis across Alzheimer’s disease and schizophrenia by integrating genetic risk and gene expression analysis. Stratified LD score regression revealed that schizophrenia genetic variants were more strongly associated with differentially expressed genes (DEGs) in AD+P than were AD variants, suggesting a shared genetic architecture that may predispose to psychotic symptoms across etiologically distinct disorders. These findings imply that while some neuronal vulnerabilities may be common to psychosis broadly, potentially enabling cross-diagnostic therapeutic strategies, other features are disease-specific and reflect the unique pathophysiological context of AD. As such, treatment development efforts may benefit from a dual approach. For instance, targeting core neuronal stress and synaptic dysregulation pathways shared across AD+P and schizophrenia, while also addressing AD+P-specific mechanisms to achieve more tailored and effective interventions.

Collectively, our findings underscore the need for a deeper mechanistic understanding of the molecular and genetic drivers of psychosis in dementia, an area that remains largely uncharted. As single-cell and multi-omic technologies continue to refine our ability to map disease-associated alterations, the field is poised to leverage this expanding molecular landscape, including datasets like ours, for targeted therapeutic development. A particularly promising avenue is the use of patient-derived iPSCs to model psychosis-associated circuit dysfunction, providing a controlled platform to dissect the genetic contributions to neuronal and glial pathology. However, given the complexity of psychiatric symptoms in dementia and our limited understanding of their genetic underpinnings, these efforts require large-scale iPSC repositories derived from diverse patient cohorts. With greater access to patient genetics, the generation of isogenic lines will allow for more precise manipulation of risk variants, accelerating the discovery of disease-modifying treatments and ultimately guiding the development of more effective therapeutics for psychosis in neurodegenerative disease.

## Methods

### Human postmortem brain samples

Postmortem brain tissue samples were obtained from the neuropathology core of the University of Pittsburgh Alzheimer’s disease research center. Subjects underwent annual neurological, neuropsychological and psychiatric evaluations as previously described ^19,23,84^. An independent committee of experienced clinicians made consensus DSM-IV diagnoses for each subject. Psychosis was defined as the presence of delusions or hallucinations at any visit. Subjects with a pre-existing psychotic disorder (e.g. schizophrenia) were excluded from the study.

Brain banking protocols were approved by the local Committee for Oversight of Research and Clinical Training Involving Decedents (CORID). At the time of autopsy, the brain was bisected and the left hemisphere fixed in formalin for subsequent diagnostic evaluation using consensus guidelines ^85–88^. The right hemisphere was dissected fresh with multiple aliquots taken from different brain regions and frozen at -80°C.

### Nuclei isolation from frozen postmortem brain tissue for snRNA-seq and snATAC-seq

We extracted nuclei from frozen postmortem brain tissue following an adapted version of the method described by Mathys et al. with some modifications. In brief, brain tissue was homogenized in 700 μL of Homogenization buffer, then filtered through a 40 μm cell strainer (Corning, NY). After adding 450 μL of Working solution, the sample was layered onto a 30%/40% OptiPrep density gradient (750 μL of 30% OptiPrep solution and 300 μL of 40% OptiPrep solution) to form a 25% OptiPrep solution. Nuclei were separated via centrifugation at 10,000g for 5 minutes at 4°C using a fixed rotor tabletop centrifuge. The nuclei pellet was collected from the 30%/40% interphase, transferred to a clean tube, washed twice with 1 mL of ice-cold PBS containing 0.04% BSA (centrifuged at 300g for 3 minutes at 4°C), and resuspended in 100 μL of PBS with 0.04% BSA. After counting, nuclei were diluted to a final concentration of 1000 nuclei per μL.

For single-nucleus RNA sequencing (snRNA-seq), the isolated nuclei were processed using the 10x Genomics Chromium Single-Cell 3′ Reagent Kits v3, targeting 5000 nuclei per brain region per individual. Library preparation was carried out following the manufacturer’s protocol, and sequencing was performed using the Illumina NovaSeq 6000 S2 system (100 cycles). For single-nucleus ATAC sequencing (snATAC-seq), the remaining nuclei were pelleted, resuspended in 1x Diluted Nuclei buffer, and adjusted to a concentration of 2500 nuclei per μL. Library construction was completed using the 10x Genomics Chromium Single-Cell ATAC Reagent Kit v1 according to the manufacturer’s instructions, with sequencing conducted on the NovaSeq 6000 S2 platform (100 cycles, Illumina).

### snRNA-seq data processing

We processed raw sequencing reads by mapping them to the human reference genome GRCh38 and quantified unique molecular identifiers (UMIs) for each gene in each cell using CellRanger v6.1.2 (10x Genomics)^89^. We then pre-processed the resulting gene-cell count matrix using Seurat v4.4.0^90^. For quality control, we retained cells with more than 500 UMIs and less than 5% mitochondrial genes, and included genes expressed in at least 50 cells for further analysis. We normalized UMI counts by total counts per cell, scaled them to 10,000 UMIs per cell, and performed log transformation. We identified the top 2000 highly variable genes for principal component analysis (PCA) and used the top 30 principal components (PCs) to generate a UMAP embedding. To correct for batch effects, we applied Harmony ^91^. Clusters were identified using a resolution of 0.5. To filter out potential doublets, we applied DoubletFinder ^92^ with default parameters and removed cells with a doublet score above 0.2. Additionally, after clustering, we excluded clusters expressing markers from two or more cell types, as they were likely doublets.

### snATAC-seq data processing

We processed snATAC-seq data by first generating raw FASTQ files for each sample through demultiplexing with cellranger-atac software(v2.0) ^93^. Next, we aligned the reads to the human reference genome (GRCh38) using “cellranger-atac count”, producing fragment files for each sample. For downstream analysis, we used ArchR (v1.0.1)^94^, removing potential doublets with the “filterDoublets” function. We retained cells with TSS enrichment >6 and fragment counts between 1,000 and 100,000 for further analysis. We performed iterative latent semantic indexing (LSI) dimension reduction and clustering using a 500 bp tile-based matrix, setting ‘iterations=4’, ‘resolution=0.2’, and ‘varFeat=50000’. We visualized cell embeddings with UMAP and generated the gene score matrix in ArchR. Cell type annotation was based on gene scores of well-established brain marker genes^95^.

### Peak calling and differential peak analysis

We identified chromatin accessible peaks using MACS2, following the ArchR pipeline^94,96^, and constructed a peak-by-cell matrix. To identify marker peaks for each cell type, we applied a Wilcoxon test in ArchR, using a p-value cutoff of <0.01 and log₂ fold change >0.5. We then annotated the marker peaks with GREAT analysis^97,98^, using the basalPlusExt model and extending 100kb upstream and downstream of each peak. For motif analysis, we expanded peaks to 2000 bp and applied Homer with default parameters ^99^.

### Co-accessibility analysis

We inferred co-accessibility between proximal and distal peaks as a proxy for peak-to-gene links, assessing correlations between peak accessibility and gene expression. We applied the following parameters: corCutOff = 0.3, FDR = 1e-4, upstream = 50,000 bp, downstream = 50,000 bp. The identified peak-gene pairs were further analyzed to link distal accessible peaks with differentially expressed genes in AD with psychosis.

### Cell proportional analysis

We performed propeller^100^ analysis to evaluate the statistical significance for compositional differences between phenotypic groups with the consideration of the confounding variables in the model including sex, age and TDP pathology. We used the *speckle* package in R and adjusted p-value < 0.05 as cutoff for significance.

### Gene Ontology enrichment analysis

We applied Enrichr in R to perform enrichment analysis for Gene Ontology biological process (adjusted p-value < 0.05 as a cutoff)^101,102^. We selected the representative terms for visualization manually based on the diverse functional categories and the number of genes involved in each term.

### Immunohistochemistry of postmortem human brains

Formalin-fixed, paraffin-embedded (FFPE) human brain tissue (prefrontal cortex) was sectioned, mounted, and processed at the neuropathology core of the University of Pittsburgh Alzheimer’s disease research center (ADRC). Slides were deparaffinized with xylene, and then rehydrated through a series of graded ethanol rinses (e.g., 100%, 95%, 70%) followed by a final rinse with 1xPBS before immunostaining. Sections were blocked for 1 h at room temperature (RT) in blocking buffer (PBS containing 5% bovine serum albumin, 1% normal donkey serum and 0.3% Triton X-100). The sections were incubated for 24 hours at 4°C with the primary antibodies in blocking buffer. NeuN (Synaptic Systems; Cat#266 004), Vglut1 (Synaptic Systems; Cat#135 302), MAP2 (BioLegend; Cat#822501), Phospho-Tau (Ser202, Thr205)(AT8) (Thermo Fisher Scientific; Cat#MN1020), Phospho-Tau (Ser396) (Thermo Fisher Scientific; Cat#44-752G). Following primary antibody incubation, the sections were washed three times for 5 min each at RT in PBS and then incubated with secondary antibodies (dilution 1:1000) for 2 h at RT. For plaque staining, Methoxy-X04 (Tocris; #4920) was added together with secondary antibodies. The slides were once again washed three times at RT in PBS and incubated in TrueBlack (Biotium; Cat#23007) for 90 seconds. After a final round of washes, the slices were mounted in Fluromount-G (SouthernBiotech; Cat#0100-01). Primary antibodies were visualized with Alexa Fluor 405, Alexa Fluor 488, Alexa Fluor 555, and Alexa Fluor 647 antibodies (Molecular Probes), and cell nuclei visualized with DAPI using a confocal microscope (LSM 900; Zeiss) with a 20× or 63× objective. For experiments quantifying fluorescent intensity, images were analyzed with Imaris imaging software (Oxford Instruments).

### In situ hybridization with RNAscope in postmortem human brains

Slides of FFPE human brain tissue corresponding to the prefrontal cortex were processed by University of Pittsburgh ADRC and shipped to ACD Biotechne for RNAscope assays and imaging. Brain sections were probed for human SYN1, STAT3, and GFAP and counterstained with DAPI. Slides were imaged with a slide scanner and files shared from ACD Biotechne were quantified with Slideviewer software from 3D Histech.

### Gene overexpression, plasmid cloning, lentivirus production

We obtained AAV-hsyn-CheRiff-eGFP from Adam Cohen through Addgene (Plasmid #51697), then subcloned it into a lentiviral vector carrying a second promoter PGK driving blasticidin resistance cassette (Addgene; #666808) to create a lentiviral construct that would allow for the generation of a stable cell line expressing neuronal-restrictived hSyn-CheRiff upon transduction and selection with blasticidin in iPSCs prior to organoid derivation. After antibiotic selection, colonies were confirmed to carry hSyn-CheRiff via PCR screening. For overexpression screen in organoids, we populated a list of 51 up-regulated DEGs from L2/L3 from the following GO terms: neuron projection morphogenesis (GO:0048812), regulation of neuron projection development (GO:0010975), positive regulation of neuron projection development (GO:0010976), neuron projection development (GO:0031175), and neuron projection organization (GO:0106027). Of these, 37 were commercially available in lentiviral vectors through the plasmid repository at DNASU (Arizona State University, dnasu.org). Received plasmids were made into lentivirus using established protocols and as reported in Victor et al., Cell Stem Cell 2022. Organoids were transduced separately with each virus or a control construct under the same conditions carrying the mammalian codon optimized variant of the blue fluorescent protein, azurite (pLv-Azurite; Addgene #36086).

### iPSC Cultures, Generation of Forebrain Organoids, and Assembloids

Human iPSC line used in this study was derived from a healthy donor (APOE3/3 genotype) and generated by the Picower Institute for Learning and Memory iPSC Facility as first described (Lin et al., 2018). iPSCs were maintained at 37°C and 5% CO2, in feeder-free conditions in mTeSR1 medium (Cat #85850; STEMCELL Technologies) on Matrigel-coated plates (Cat # 354277; Corning; hESC-Qualified Matrix). iPSCs were passaged at 60–80% confluence using ReLeSR (Cat# 05872; STEMCELL Technologies) and reseeded 1:6 onto Matrigel-coated plates. Dorsal forebrain organoids were generated using a previously established protocol (Sloan et al., 2018) with an adapted iPS seeding strategy (Marton et al., 2019), as described previously in Victor et al., Cell Stem Cell 2022. After 60 days of culture, organoids were transduced with lentivirus or AAVs and cultured separately for an additional 2 weeks. Next, pairs of organoids with desired infected constructs were moved to microcentrifuge tubes and allowed to fuse for 24 hours to generate assembloids. Following fusion, assembloids were cultured separately in untreated glass bottom 24-well microplates (Cat #82421; Ibidi) and fed every 3-4 days with experiments starting approximately at 90 days in vitro.

### Calcium Reporter, Imaging and Optogenetic Stimulation

E.SV40 (Cat#100843-AAV1; Addgene) while temporarily plated in 24-well plates (3-4 organoids per well) with 20ul of viral prep (100 µL at titer ≥ 1×10¹³ vg/mL). After overnight incubation with virus, organoids were returned to ultra-low attachment 10cm dishes. Typically within 12-14 days calcium dynamics were easily detectable. For experiments with AAV Axonal GCaMP6s, a similar protocol was followed but with pAAV-hSynapsin1-axon-GCaMP6s (Plasmid #111262). For calcium imaging, live-imaging was performed with Zeiss LSM900 equipped with a heated chamber kept at 37 C with humidity and CO2 control. Images were processed and analyzed as described previously (Victor et al., Cell Stem Cell 2022). Calcium traces from motion-corrected time series were manually segmented on ImageJ into individual cells based on threshold intensity, variance, and upper and lower limits for cell size. Image segmentation results were separately inspected for quality control. Fluorescence signal time series (DF/F: change in fluorescence divided by baseline fluorescence) were calculated for each segment. The baseline fluorescence for each cell was determined as the minimum fluorescence signal in the baseline recording epoch. For GCaMP-tagged neurons, The onset of a calcium transient occurred when DF/F exceeded two standard deviations above the baseline fluorescence. The termination of a calcium transient was identified as occurring when DF/F fell below 0.5 standard deviations above the baseline fluorescence. We defined a multicellular ensemble events when the number of simultaneously active cells exceeded 60% of all cells. For CheRiff-GCaMP6s assembloid screen, optogenetic stimulation and calcium imaging were coupled and acquired using low magnification 2.5x objective to scan the entire assembloid structure. For experiments with chronic optogenetic stimulation, custom LED tissue culture plates were outfitted under the control of an Arduino board and stimulation carried out for 200ms at 10Hz every 2 minutes for 48 hours. Heatmaps were generated using GraphPad Prism (GraphPad Software).

### Patch-Clamp Electrophysiology

Whole-cell patch-clamp recordings of neurons were performed at 6 to 8 weeks after NGN2 induction from hSyn-CheRiff iPSC line. Intracellular recordings were performed at room temperature using an Axon CV-7B headstage, Multiclamp 700B amplifier, and Digidata 1440A digitizer (Molecular Devices). Electrode pipettes were pulled from borosilicate glass on a Model P-97 Flaming/Brown micropipette puller (Sutter Instrument) and ranged between 4–7 MΩ resistance. Intrinsic neuronal properties were studied using the following solutions (in mM): Extracellular: 125 NaCl, 2.5 KCl, 1.2 NaH2PO42H2O, 1.2 MgCl26H2O, 2.4 CaCl22H2O, 26 NaHCO3, 11 glucose (pH 7.4). Intracellular: 135 K-gluconate, 5 KCl, 2 MgCl26H2O, 10 HEPES, 2 Mg-ATP, 0.2 Na2GTP (pH 7.2). Membrane potentials were typically kept at -70 mV. In current-clamp mode, action potentials were elicited by photostimulation with blue light. Data was first collected and analyzed using pCLAMP 11 software (Molecular Devices).

### MEA Recordings

Intact organoids were plated onto Poly-D-Lysine (Cat#P6407-10X5MG; Sigma-Aldrich) coated wells of a CytoView MEA 48-well plate (Cat#M768-tMEA-48B; Axion BioSystems) and covered in a Matrigel droplet (Cat # 354277; Corning) to anchor the organoid. After 15–30 min in 37 C, droplets were flooded with warm BrainPhys Medium (Cat#05790; STEMCELL Technologies) and allowed to recover for at least four weeks before recording sessions. A recording session preceded the 5ng/ml IL-6 treatment, denoted as baseline recording. For optogenetic stimulation, organoids were pulsed continuously at 6Hz with blue light. All extracellular recordings were performed using the Axion Maestro Pro MEA system (Axion Biosystems). Spontaneous neural activity was recorded for 15 min at a sampling rate of 12.5 kHz, and an adaptive threshold set at 5.5 times the standard deviation of baseline noise was used for spike detection. Electrodes were defined as active if neuronal firing occurred at a minimal rate of 5 spikes/min. For MEA data analysis, only wells containing a minimum of 3 active electrodes were included. Neuronal firing metrics were exported as the averages from each well from Axion Biosystems’ Neural Metrics Tool and plotted with Prism GraphPad (GraphPad Software). For optogenetic stimulation, custom blue LED panels were adapted to fit the Axion Maestro Pro and delivered under the control of an Arduino board at 6Hz.

### Gene Expression analysis with qPCR

RNA extraction for qPCR analysis was performed with RNeasy Plus Mini Kit (Cat# Cat#74134; Qiagen), and reverse transcription was performed with RNA to cDNA EcoDry Premix (Cat# Cat#639549; Takara) according to the manufacturer’s instructions. Gene expression was analyzed with Real-Time PCR (CFX96; Bio-Rad) and SsoFast EvaGreen Supermix (Cat#1725202; Bio-Rad). Expression data was normalized to house-keeping gene GAPDH using the 2 DDCT relative quantification method. CDK5R1 expression was assessed with primers (5’ - > 3’): Forward - AGAACAGCAAGAACGCCAAG and Reverse - CGGCCACGATTCTCTTCCA. Housekeeping primers for GAPDH (5’ -> 3’): Forward - AGAAGGCTGGGGCTCATTTG and Reverse - AGGGGCCATCCACAGTCTTC.

### Identification of differentially expressed genes between AD with and without psychosis

We applied two methods (one single-cell based method, MAST, and a single-cell based method edgeR) to detect the differentially expressed genes for each cell type between AD with and without psychosis individuals per brain region. We controlled the covariates for both methods, including percentage of mitochondrial genes, percentage of ribosomal genes, age, sex, PMI, neocortical Lewy body, and TDP pathology. The genes with FDR <= 0.05 for MAST and p-value <= 0.05 for edgeR were selected for further analysis. We found the DEGs were highly consistent between two methods.

### Prediction of upstream transcription factors

We inferred the upstream transcription factors of genes (marker genes or DEGs) using Enrichr in R for the enrichment-based analysis based on three libraries including TRANSFAC and JASPAR, ChEA, and ENCODE TF ChIP-seq data ^66,103,104^. We used adjusted p-value <0.05 as a cut-off to select the significant TFs. For motif-based analysis, we linked the DEGs to distal peaks using the peak-to-gene links represented by co-accessible peaks and then performed motif analysis by expanding the peaks to 2000bp with the default parameters in Homer ^99^. We kept the TFs with detected expression (average expression > 1 after log2 transformation) in relevant cell types for further analysis.

### LDSC Heritability Enrichment Analysis

We applied Stratified Linkage Disequilibrium Score Regression (S-LDSC)^74^ to evaluate the heritability enrichment of genetic risk for Schizophrenia and AD within DEGs related accessible chromatin regions. To focus on functionally relevant elements, we first identified distal accessible peaks linked to DEGs in AD+P individuals by leveraging co-accessibility between promoter and distal regulatory regions. For these DEG-linked peaks, we generated binary annotation files indicating whether each SNP overlapped with an accessible region. To account for genetic variants in linkage disequilibrium (LD) and address the sparsity of variant annotations, we extended annotated regions by 1000 bp to both directions. We formatted these annotations according to the standard S-LDSC pipeline and computed LD scores using the 1000 Genomes Project Phase 3 European reference panel. We then estimated partitioned heritability with publicly available Schizophrenia and AD GWAS summary statistics^105,106^, which include SNP-level effect sizes and standard errors. We defined enrichment as the proportion of heritability explained by SNPs within accessible peaks relative to the proportion of SNPs in those peaks. A significant enrichment suggests that disease-associated variants are preferentially located within regulatory elements specific to the cell types under investigation.

### Data availability statement

The dataset generated in this study will be made publicly available upon publication. Additional supporting datasets are available from the corresponding author upon reasonable request. All code used for data processing and analysis will be released at https://github.com/nasunmit/ at the time of publication.

For peer review purposes, the data has been deposited on GEO. The following secure token has been created to allow review of record GSE296869 while it remains in private status:

To review GEO accession GSE296869: Go to https://www.ncbi.nlm.nih.gov/geo/query/acc.cgi?acc=GSE296869 Enter token atkvaqkqhhqhhwv into the box

## Supplementary Figure Legend

**Figure S1. Histology for Neuropathological Changes in AD-P and AD+P.** An additional cohort of postmortem human brain tissue from 20 subjects (n= 10 AD-P and 10 AD+P). Error bar represents the SEM. Two-way ANOVA with Bonferroni’s multiple comparison test.

**Figure S2. Cell Fraction Analysis for Anterior Hippocampus in AD-P and AD+P.** Propeller analysis was used to determine statistical significance. Error bar represents the SEM. ns= not significant. n= 24 AD+P and 24 AD-P postmortem human brain tissue. AH: Anterior Hippocampus.

**Figure S3.Transcriptional Changes in AD with Psychosis. A-B.** Scatter plot to show the consistency of transcriptional changes identified by two methods in SF (**A**) and AH (**B**). The X-axis represents the coefficient calculated by MAST, while the Y-axis represents the log transformed fold change calculated by edgeR.

**Figure S4. Gene Ontology (GO) for Transcriptional changes in AD with psychosis.** Representative Gene Ontology from significantly enriched terms for five cell types.

**Figure S5. Compensatory Mechanism Evoked by Surviving Excitatory Neurons and Correlates of Metabolic Stress in PV+ Interneurons. A - B.** Histology of PFC human brain sections with H&E (hematoxylin and eosin) staining and fluorescent in situ RNA hybridization using an RNAscope probe for synapsin 1 (SYN1) from AD+P (n=3) and AD-P (n=3) subjects. **C.** Quantification of RNAscope dots per nucleus for a given SYN1-positive neuron (comprising at least 3 SYN1 dots) increased in AD+P across cortical layers. *p-value <0.05, multiple t-tests adjusted for false discovery rate. And in Layers L2/L3. **D.** Higher magnification insets of layers L1, L2 and L3 with SYN1 probe in green and DAPI in blue. **E.** Percentage of cells SYN1-positive ( at least 3 SYN1 dots) in cortical layers L2 and L3 were significantly reduced in AD+P. (n=averages of 3 subjects per group). *p-value <0.05, **p-value <0.01, Student’s t-test. **F.** Cell fraction analysis from PFC snRNA-seq of inhibitory interneurons. n= 24 subjects per group of snRNA-seq. Propeller analysis to determine statistical significance. **G.** Top GO terms from AD+P DEGs of Parvalbumin (PV)-positive cells. Error bars represent S.E.M.

**Figure S6. CAMK2B Overexpression Renders Forebrain Organoids Vulnerable to Excitotoxicity. A.** Custom LED Tissue Culture Plate. **B-C**. Diagram of experimental design **D**. Stimulation protocol. **E**. Control organoids were tagged with pLenti CMV-EBFP. **F**. Organoids tagged with AAV hSYN-RFP before (baseline) and after stimulation (post-stim) and quantified (**G**). Paired t-test; **p-value<0.01, ns= not significant. n=10 control and 10 CAMK2B overpressing organoids).

**Figure S7. Epigenetic Regulation of Cell-type-specific Gene Expression Changes. A.** Dotplot shows the gene activity score of well-known marker genes for major cell types in the human brain in snATAC-seq data. The color represents the activity level and the size of the dot represents the percentage of cells with activity. **B**. The composition of major cell types in snATAC-seq data in two brain regions, AH and SF. **C.** The comparison of cell fractions captured by snATAC-seq and snRNA-seq data in two brain regions. **D-E**. UMAPs to show the integrative analysis of snATAC-seq and snRNA-seq data in two brain regions. The UMAP on the left represents the joint map, and the right panel represents the nuclei from single modality. The nuclei were colored by major cell types. **F.** The number of peaks per cell type and union set, colored by their genomic locations (distal, exonic, intronic or promoter regions). **G**. The heamap to show the links between distal peak to promoter peaks, representing the co-accessbility between distal enhancer and promoter. Each row represents one link and each column represents randomly selected nuclei (500 in total for visualization). The left panel represents the activity in snATAC-seq, and the right plan represents the gene expression. Z-score values were used. Most of the links were cell-type specific.

**Supplemental Figure S8. Limited concordance between canonical AD and AD+P transcriptional signatures across major brain cell types.** Scatter plots show the relationship between differential gene expression effect sizes in Alzheimer’s disease (AD, x-axis), re-analyzed from Mathys et al. using the same differential expression framework applied in this study, and Alzheimer’s disease with psychosis (AD+P, y-axis). Differential expression was computed relative to the appropriate control group for each dataset across all major brain cell classes, including excitatory neurons (ExN), inhibitory neurons (InN), astrocytes (Ast), oligodendrocytes (Oli), oligodendrocyte precursor cells (Opc), and microglia (Mic). Each point represents a gene tested for differential expression. Marginal histograms depict the distribution of effect sizes along each axis. This analysis illustrates both shared and divergent transcriptional responses between AD and AD+P across neuronal and glial populations.

**Supplemental Figure S9. Cross-cell-type overlap of differential gene expression programs in AD and AD+P.** Heatmap depicts the degree of overlap between differentially expressed gene sets identified in Alzheimer’s disease without psychosis (AD) and Alzheimer’s disease with psychosis (AD+P) across major brain cell types, including excitatory neurons (ExN), inhibitory neurons (InN), astrocytes (Ast), oligodendrocyte precursor cells (Opc), oligodendrocytes (Oli), and microglia (Mic). Rows correspond to AD gene sets and columns to AD+P gene sets, each separated into up- and down-regulated categories. Cell values indicate the magnitude of gene overlap or enrichment between each gene set pair, as displayed by the color scale. This analysis highlights shared and distinct transcriptional programs across neuronal and glial populations in AD and AD+P.

**Figure S10. Cell-type-specific Gene Expression Changes and their Association with Schizophrenia Genetics. H.** AD+P DEGs linked chromatin accessible peaks show the less enrichment of genetic variants of AD using LDSC analysis. The barplot shows the enrichment score, where pink ones represent the significance. The -log10 of p-value is shown by the blue point. The red dash line represents the stringent threshold with p-value < 0.01. **I-J**. The UpSet plot shows the overlap of DEGs in inhibitory neurons (**I**) and glial cell types (**J**) between Alzheimer’s Disease (AD), Schizophrenia (Sz) and AD+P (ADP).

## Supplementary Tables and Movies

**Table S1. Patient Manifest for snRNA-seq and snATAC-seq.**

**Table S2. Patient Manifest for Histology Experiments.**

**Table S3. Prefrontal Cortex (PFC) DEGs.**

**Table S4. Anterior Hippocampus (AH) DEGs.**

**Table S5. PFC Non-Neuronal GO Terms.**

**Table S6. GREAT Analysis. Table S7. Peak-to-Gene Links.**

**Movie S1. Axonal GCaMP at Assembloid Junction Line.**

**Movie S2. Representative CheRiff-GCaMP Assembloid.**

**Movie S3. Functional Screen for L2/L3 DEGs on Assembloids**

## Acknowledgements

We thank the study participants and staff of the Alzheimer Disease Research Center at the University of Pittsburgh and the brain banking teams and histology staff of the neurodegenerative brain bank at the University of Pittsburgh. The Tsai lab research was made possible through the generous support of the Robert A. and Renee E. Belfer Family Foundation, The Freedom Together Foundation, Eduardo Eurnekian, Lester A. Gimpelson, the Donald A. Mattes (1967) Memorial Neuroscience Research Expendable Fund and Glenda G. Mattes and Steve Corbin, David Emmes, the J. Crayton Pruitt Foundation, and Joseph P. DiSabato and Nancy E. Sakamoto. This work was also supported in part by NIH grants: R01AG062335 (M.K. and L.-H.T.), R01AG081017 (M.K.), R01MH109978 (M.K.), U01MH119509 (M.K.), R01AG074003 (M.K. and L.-H.T.), U01NS110453 (M.K. and L.-H.T.), R01AG058002 (M.K. and L.-H.T.), RF1AG062377 (L.-H.T.), RF1AG054012 (L.-H.T. and M.K.), R01AG067151 (M.K.), UH3NS115064 (L.-H.T. and M.K.), RF1NS129032 (M.K.), R01NS127187 (M.K.), U01AG077227 (L.-H.T.), U01HG012009 (M.K.), U01DA053631 (M.K.), R01DA054584 (M.K.), R01HG008155 (M.K.), H116046 (J.K.K., and R.A.S.), P30-AG066468 (J.K.K., and R.A.S.) and grants from the Cure Alzheimer’s Fund CIRCUITS consortium (M.K. and L.-H.T), The Biswas Family Foundation and the Milken Institute (M.K.), and the Alana Foundation (M.K.). M.B.V. is supported by the Howard Hughes Medical Institute Hanna H. Gray Fellowship.

## Author contributions

M.B.V, N.S., R.A.S., L.-H.T., and M.K. conceived and designed the study; R.A.S., M.K. and L.-H.T. supervised the study; J.K.K. supervised histological experiments, guided data interpretation, and collaborated in study design; N.S. developed the computational framework and conducted data analysis with assistance from Y.T.; M.B.V., N.L., A.N.S., and S.P. performed in vitro experiments and analyzed results; K.G., H.M., X.J., L.-L.H. and A.P.N. performed snRNA-seq and snATAC-seq profiling; L.L. performed electrophysiology analysis; N.S., J.K.K. and M.B.V wrote methods; R.A.S. and J.K.K. provided postmortem samples and scientific input; and M.B.V, N.S., R.A.S., L.-H.T. and M.K. wrote and revised the manuscript.

## References

1. Lively, S. T. Activities of Daily Living (ADL): Cultural Differences, Impacts of Disease and Long-Term Health Effects. (Nova Science Publishers, 2015).

2. The spectrum of behavioral changes in Alzheimer’s disease. *Neurology* https://www.neurology.org/doi/10.1212/WNL.46.1.130.

3. Knopman, D. S. et al. Alzheimer disease. Nature Reviews Disease Primers 7, 1–21 (2021).

4. Schuff, N. et al. MRI of hippocampal volume loss in early Alzheimer’s disease in relation to ApoE genotype and biomarkers. Brain 132, 1067–1077 (2009).

5. Salat, D. H., Kaye, J. A. & Janowsky, J. S. Prefrontal gray and white matter volumes in healthy aging and Alzheimer disease. Arch Neurol 56, 338–344 (1999).

6. Canter, R. G., Penney, J. & Tsai, L.-H. The road to restoring neural circuits for the treatment of Alzheimer’s disease. Nature 539, 187–196 (2016).

7. Xu Lou, I., Chen, J., Ali, K., Shaikh, A. L. & Chen, Q. Mapping new pharmacological interventions for cognitive function in Alzheimer’s disease: a systematic review of randomized clinical trials. Front Pharmacol 14, 1190604 (2023).

8. Murray, P. S., Kumar, S., Demichele-Sweet, M. A. A. & Sweet, R. A. Psychosis in Alzheimer’s disease. Biol Psychiatry 75, 542–552 (2014).

9. Ismail, Z. et al. Psychosis in Alzheimer disease - mechanisms, genetics and therapeutic opportunities. Nat Rev Neurol 18, 131–144 (2022).

10. American Psychiatric Association. The American Psychiatric Association Practice Guideline on the Use of Antipsychotics to Treat Agitation or Psychosis in Patients With Dementia. (American Psychiatric Pub, 2016).

11. Narang, P. et al. Antipsychotic drugs: sudden cardiac death among elderly patients. Psychiatry (Edgmont) 7, 25–29 (2010).

12. Schneider, L. S. & Dagerman, K. S. Psychosis of Alzheimer’s disease: clinical characteristics and history. J Psychiatr Res 38, 105–111 (2004).

13. DeMichele-Sweet, M. A. A. et al. Genome-wide association identifies the first risk loci for psychosis in Alzheimer disease. Mol Psychiatry 26, 5797–5811 (2021).

14. DeMichele-Sweet, M. A. A. et al. Genetic risk for schizophrenia and psychosis in Alzheimer disease. Mol Psychiatry 23, 963–972 (2018).

15. Shah, C., DeMichele-Sweet, M. A. A. & Sweet, R. A. Genetics of psychosis of Alzheimer disease. Am J Med Genet B Neuropsychiatr Genet 174, 27–35 (2017).

16. Antonsdottir, I. M. et al. Genetic associations with psychosis and affective disturbance in Alzheimer’s disease. Alzheimers Dement (N Y) 10, e12472 (2024).

17. Jauhar, S. et al. The relationship between cortical glutamate and striatal dopamine in first-episode psychosis: a cross-sectional multimodal PET and magnetic resonance spectroscopy imaging study. Lancet Psychiatry 5, 816–823 (2018).

18. Basavaraju, R. et al. Hippocampal Glutamate and Positive Symptom Severity in Clinical High Risk for Psychosis. JAMA Psychiatry 79, 178–179 (2022).

19. DeChellis-Marks, M. R. et al. Psychosis in Alzheimer’s Disease Is Associated With Increased Excitatory Neuron Vulnerability and Post-transcriptional Mechanisms Altering Synaptic Protein Levels. Front Neurol 13, 778419 (2022).

20. Kouhsar, M. et al. A brain DNA co-methylation network analysis of psychosis in Alzheimer’s disease. Alzheimers Dement 21, e14501 (2025).

21. Rikandi, E. et al. Functional network connectivity and topology during naturalistic stimulus is altered in first-episode psychosis. Schizophr Res 241, 83–91 (2022).

22. Song, X. et al. Bioenergetics and abnormal functional connectivity in psychotic disorders. Mol Psychiatry 26, 2483–2492 (2021).

23. Murray, P. S. et al. Hyperphosphorylated tau is elevated in Alzheimer’s disease with psychosis. J Alzheimers Dis 39, 759–773 (2014).

24. Almeida, F. C. et al. Psychosis in Alzheimer’s disease is associated with specific changes in brain MRI volume, cognition and neuropathology. Neurobiol Aging 138, 10–18 (2024).

25. Gomar, J. J. et al. Increased retention of tau PET ligand [F]-AV1451 in Alzheimer’s Disease Psychosis. Transl Psychiatry 12, 82 (2022).

26. DeTure, M. A. & Dickson, D. W. The neuropathological diagnosis of Alzheimer’s disease. Molecular Neurodegeneration 14, 1–18 (2019).

27. Hedegaard, C. et al. Porcine synapsin 1: SYN1 gene analysis and functional characterization of the promoter. FEBS Open Bio 3, 411–420 (2013).

28. Pickel, V. M. & Segal, M. The Synapse: Structure and Function. (Elsevier, 2013).

29. Zhang, Z. & Sun, Q.-Q. The balance between excitation and inhibition and functional sensory processing in the somatosensory cortex. Int Rev Neurobiol 97, 305–333 (2011).

30. Liu, Y. et al. A Selective Review of the Excitatory-Inhibitory Imbalance in Schizophrenia: Underlying Biology, Genetics, Microcircuits, and Symptoms. Front Cell Dev Biol 9, 664535 (2021).

31. Beasley, C. L. & Reynolds, G. P. Parvalbumin-immunoreactive neurons are reduced in the prefrontal cortex of schizophrenics. Schizophr Res 24, 349–355 (1997).

32. Beasley, C. L., Zhang, Z. J., Patten, I. & Reynolds, G. P. Selective deficits in prefrontal cortical GABAergic neurons in schizophrenia defined by the presence of calcium-binding proteins. Biol Psychiatry 52, 708–715 (2002).

33. Bitanihirwe, B. K. Y. & Woo, T.-U. W. Transcriptional dysregulation of γ-aminobutyric acid transporter in parvalbumin-containing inhibitory neurons in the prefrontal cortex in schizophrenia. Psychiatry Res 220, 1155–1159 (2014).

34. Agetsuma, M., Hamm, J. P., Tao, K., Fujisawa, S. & Yuste, R. Parvalbumin-Positive Interneurons Regulate Neuronal Ensembles in Visual Cortex. Cereb Cortex 28, 1831–1845 (2018).

35. Hu, H., Gan, J. & Jonas, P. Interneurons. Fast-spiking, parvalbumin+ GABAergic interneurons: from cellular design to microcircuit function. Science 345, 1255263 (2014).

36. Hensch, T. K. & Fagiolini, M. Excitatory-Inhibitory Balance: Synapses, Circuits, Systems. (Springer Science & Business Media, 2012).

37. Singer, W., Sejnowski, T. J. & Rakic, P. The Neocortex. (MIT Press, 2019).

38. Gerfen, C. R., Economo, M. N. & Chandrashekar, J. Long distance projections of cortical pyramidal neurons. J Neurosci Res 96, 1467–1475 (2018).

39. Blanchard, J. W., Victor, M. B. & Tsai, L.-H. Dissecting the complexities of Alzheimer disease with in vitro models of the human brain. Nat Rev Neurol 18, 25–39 (2022).

40. Sloan, S. A., Andersen, J., Pașca, A. M., Birey, F. & Pașca, S. P. Generation and assembly of human brain region-specific three-dimensional cultures. Nat Protoc 13, 2062–2085 (2018).

41. Hayward, R. F., Brooks, F. P., 3rd, Yang, S., Gao, S. & Cohen, A. E. Diminishing neuronal acidification by channelrhodopsins with low proton conduction. Elife 12, (2023).

42. Broussard, G. J. et al. In vivo measurement of afferent activity with axon-specific calcium imaging. Nat Neurosci 21, 1272–1280 (2018).

43. Nicole, O. & Pacary, E. CaMKIIβ in Neuronal Development and Plasticity: An Emerging Candidate in Brain Diseases. Int J Mol Sci 21, (2020).

44. Vest, R. S., O’Leary, H., Coultrap, S. J., Kindy, M. S. & Bayer, K. U. Effective post-insult neuroprotection by a novel Ca(2+)/ calmodulin-dependent protein kinase II (CaMKII) inhibitor. J Biol Chem 285, 20675–20682 (2010).

45. Tsai, L. H., Delalle, I., Caviness, V. S., Jr, Chae, T. & Harlow, E. p35 is a neural-specific regulatory subunit of cyclin-dependent kinase 5. Nature 371, 419–423 (1994).

46. Chae, T. et al. Mice lacking p35, a neuronal specific activator of Cdk5, display cortical lamination defects, seizures, and adult lethality. Neuron 18, 29–42 (1997).

47. Noble, W. et al. Cdk5 is a key factor in tau aggregation and tangle formation in vivo. Neuron 38, 555–565 (2003).

48. Lee, M. S. et al. Neurotoxicity induces cleavage of p35 to p25 by calpain. Nature 405, 360–364 (2000).

49. Seo, J. et al. Activity-dependent p25 generation regulates synaptic plasticity and Aβ-induced cognitive impairment. Cell 157, 486–498 (2014).

50. Patrick, G. N. et al. Conversion of p35 to p25 deregulates Cdk5 activity and promotes neurodegeneration. Nature 402, 615–622 (1999).

51. Pao, P.-C. & Tsai, L.-H. Three decades of Cdk5. J Biomed Sci 28, 79 (2021).

52. Tsai, L. H., Takahashi, T., Caviness, V. S., Jr & Harlow, E. Activity and expression pattern of cyclin-dependent kinase 5 in the embryonic mouse nervous system. Development 119, 1029–1040 (1993).

53. Kimura, T., Ishiguro, K. & Hisanaga, S.-I. Physiological and pathological phosphorylation of tau by Cdk5. Front Mol Neurosci 7, 65 (2014).

54. Nabizadeh, F. et al. Neuroimaging Findings of Psychosis in Alzheimer’s Disease: A Systematic Review. Brain Behav 15, e70205 (2025).

55. Sun, N. et al. Human microglial state dynamics in Alzheimer’s disease progression. Cell 186, 4386–4403.e29 (2023).

56. Keren-Shaul, H. et al. A Unique Microglia Type Associated with Restricting Development of Alzheimer’s Disease. Cell 169, 1276–1290.e17 (2017).

57. Mathys, H. et al. Temporal Tracking of Microglia Activation in Neurodegeneration at Single-Cell Resolution. Cell Rep 21, 366–380 (2017).

58. Olah, M. et al. Single cell RNA sequencing of human microglia uncovers a subset associated with Alzheimer’s disease. Nat Commun 11, 6129 (2020).

59. Tuddenham, J. F. et al. A cross-disease resource of living human microglia identifies disease-enriched subsets and tool compounds recapitulating microglial states. Nat Neurosci 27, 2521–2537 (2024).

60. Walker, D. G. et al. Patterns of Expression of Purinergic Receptor P2RY12, a Putative Marker for Non-Activated Microglia, in Aged and Alzheimer’s Disease Brains. Int J Mol Sci 21, (2020).

61. Tsai, A. P. et al. Genetic variants of phospholipase C-γ2 alter the phenotype and function of microglia and confer differential risk for Alzheimer’s disease. Immunity 56, 2121–2136.e6 (2023).

62. Ghoreschi, K., Laurence, A. & O’Shea, J. J. Janus kinases in immune cell signaling. Immunol Rev 228, 273–287 (2009).

63. López-Domínguez, J. A. et al. transcript variant 2 is a marker of aging and cellular senescence. Aging (Albany NY) 13, 13380–13392 (2021).

64. Karabag, D. et al. Characterizing microglial senescence: Tau as a key player. J Neurochem 166, 517–533 (2023).

65. Carling, G. K. et al. Alzheimer’s disease-linked risk alleles elevate microglial cGAS-associated senescence and neurodegeneration in a tauopathy model. Neuron 112, 3877–3896.e8 (2024).

66. Xie, Z. et al. Gene Set Knowledge Discovery with Enrichr. Curr Protoc 1, e90 (2021).

67. Huang, B., Lang, X. & Li, X. The role of IL-6/JAK2/STAT3 signaling pathway in cancers. Front Oncol 12, 1023177 (2022).

68. Puel, A. & Casanova, J.-L. The nature of human IL-6. J Exp Med 216, 1969–1971 (2019).

69. Sun, L. et al. Neuroprotection by IFN-γ via astrocyte-secreted IL-6 in acute neuroinflammation. Oncotarget 8, 40065–40078 (2017).

70. Lyra E Silva, N. M., et al. Pro-inflammatory interleukin-6 signaling links cognitive impairments and peripheral metabolic alterations in Alzheimer’s disease. Transl Psychiatry 11, 251 (2021).

71. Quintanilla, R. A., Orellana, D. I., González-Billault, C. & Maccioni, R. B. Interleukin-6 induces Alzheimer-type phosphorylation of tau protein by deregulating the cdk5/p35 pathway. Exp Cell Res 295, 245–257 (2004).

72. Xiong, X. et al. Epigenomic dissection of Alzheimer’s disease pinpoints causal variants and reveals epigenome erosion. Cell 186, 4422–4437.e21 (2023).

73. Mathys, H. et al. Single-cell atlas reveals correlates of high cognitive function, dementia, and resilience to Alzheimer’s disease pathology. Cell 186, 4365–4385.e27 (2023).

74. Finucane, H. K. et al. Partitioning heritability by functional annotation using genome-wide association summary statistics. Nat Genet 47, 1228–1235 (2015).

75. Pooler, A. M., Noble, W. & Hanger, D. P. A role for tau at the synapse in Alzheimer’s disease pathogenesis. Neuropharmacology 76 **Pt A**, 1–8 (2014).

76. Steullet, P. et al. Oxidative stress-driven parvalbumin interneuron impairment as a common mechanism in models of schizophrenia. Mol Psychiatry 22, 936–943 (2017).

77. Wu, W. et al. Microglial depletion aggravates the severity of acute and chronic seizures in mice. Brain Behav Immun 89, 245–255 (2020).

78. Kinoshita, S. & Koyama, R. Pro- and anti-epileptic roles of microglia. Neural Regen Res 16, 1369–1371 (2021).

79. Badimon, A. et al. Negative feedback control of neuronal activity by microglia. Nature 586, 417–423 (2020).

80. Ku, T. S., et al. Proteomic Analysis in Alzheimer’s Disease with Psychosis Reveals Separate Molecular Signatures for Core AD Proteinopathy and Postsynaptic Density Disruption. bioRxiv (2025) doi:10.1101/2025.11.26.690872.

81. Seo, J. et al. Inhibition of p25/Cdk5 Attenuates Tauopathy in Mouse and iPSC Models of Frontotemporal Dementia. J Neurosci 37, 9917–9924 (2017).

82. Pao, P.-C. et al. A Cdk5-derived peptide inhibits Cdk5/p25 activity and improves neurodegenerative phenotypes. Proc Natl Acad Sci U S A 120, e2217864120 (2023).

83. Zheng, Y.-L., Li, B.-S., Amin, N. D., Albers, W. & Pant, H. C. A peptide derived from cyclin-dependent kinase activator (p35) specifically inhibits Cdk5 activity and phosphorylation of tau protein in transfected cells. Eur J Biochem 269, 4427–4434 (2002).

84. Krivinko, J. M. et al. Synaptic Proteome Compensation and Resilience to Psychosis in Alzheimer’s Disease. Am J Psychiatry 175, 999–1009 (2018).

85. Montine, T. J. et al. National Institute on Aging-Alzheimer’s Association guidelines for the neuropathologic assessment of Alzheimer’s disease: a practical approach. Acta Neuropathol 123, 1–11 (2012).

86. Hyman, B. T. et al. National Institute on Aging-Alzheimer’s Association guidelines for the neuropathologic assessment of Alzheimer’s disease. Alzheimers Dement 8, 1–13 (2012).

87. McKeith, I. G. et al. Diagnosis and management of dementia with Lewy bodies: Fourth consensus report of the DLB Consortium. Neurology 89, 88–100 (2017).

88. Nelson, P. T. et al. Limbic-predominant age-related TDP-43 encephalopathy (LATE): consensus working group report. Brain 142, 1503–1527 (2019).

89. Zheng, G. X. Y. et al. Massively parallel digital transcriptional profiling of single cells. Nat. Commun. 8, 14049 (2017).

90. Hao, Y. et al. Integrated analysis of multimodal single-cell data. Cell 184, 3573–3587.e29 (2021).

91. Korsunsky, I. et al. Fast, sensitive and accurate integration of single-cell data with Harmony. Nat. Methods 16, 1289–1296 (2019).

92. McGinnis, C. S., Murrow, L. M. & Gartner, Z. J. DoubletFinder: Doublet Detection in Single-Cell RNA Sequencing Data Using Artificial Nearest Neighbors. Cell Syst 8, 329–337.e4 (2019).

93. Satpathy, A. T. et al. Massively parallel single-cell chromatin landscapes of human immune cell development and intratumoral T cell exhaustion. Nat. Biotechnol. 37, 925–936 (2019).

94. Granja, J. M. et al. ArchR is a scalable software package for integrative single-cell chromatin accessibility analysis. Nat. Genet. 53, 403–411 (2021).

95. Mathys, H. et al. Single-cell transcriptomic analysis of Alzheimer’s disease. Nature 570, 332–337 (2019).

96. Zhang, Y. et al. Model-based analysis of ChIP-Seq (MACS). Genome Biol. 9, R137 (2008).

97. McLean, C. Y. et al. GREAT improves functional interpretation of cis-regulatory regions. Nat. Biotechnol. 28, 495–501 (2010).

98. Tanigawa, Y., Dyer, E. S. & Bejerano, G. WhichTF is functionally important in your open chromatin data? PLoS Comput Biol 18, e1010378 (2022).

99. Heinz, S. et al. Simple combinations of lineage-determining transcription factors prime cis-regulatory elements required for macrophage and B cell identities. Mol. Cell 38, 576–589 (2010).

100. Phipson, B. et al. propeller: testing for differences in cell type proportions in single cell data. Bioinformatics 38, 4720–4726 (2022).

101. Ashburner, M. et al. Gene Ontology: tool for the unification of biology. Nat. Genet. 25, 25–29 (2000).

102. The Gene Ontology Consortium. The Gene Ontology Resource: 20 years and still GOing strong. Nucleic Acids Res. 47, D330–D338 (2019).

103. Chen, E. Y. et al. Enrichr: interactive and collaborative HTML5 gene list enrichment analysis tool. BMC Bioinformatics 14, 128 (2013).

104. Kuleshov, M. V. et al. Enrichr: a comprehensive gene set enrichment analysis web server 2016 update. Nucleic Acids Res. 44, W90–7 (2016).

105. Backman, J. D. et al. Exome sequencing and analysis of 454,787 UK Biobank participants. Nature 599, 628–634 (2021).

106. Bellenguez, C. et al. New insights into the genetic etiology of Alzheimer’s disease and related dementias. Nat Genet 54, 412–436 (2022).

