## Supplemental Figures S1-S10 for "Compensatory Plasticity Defines a Vulnerable Neuronal State in Alzheimer’s Disease with Psychosis"

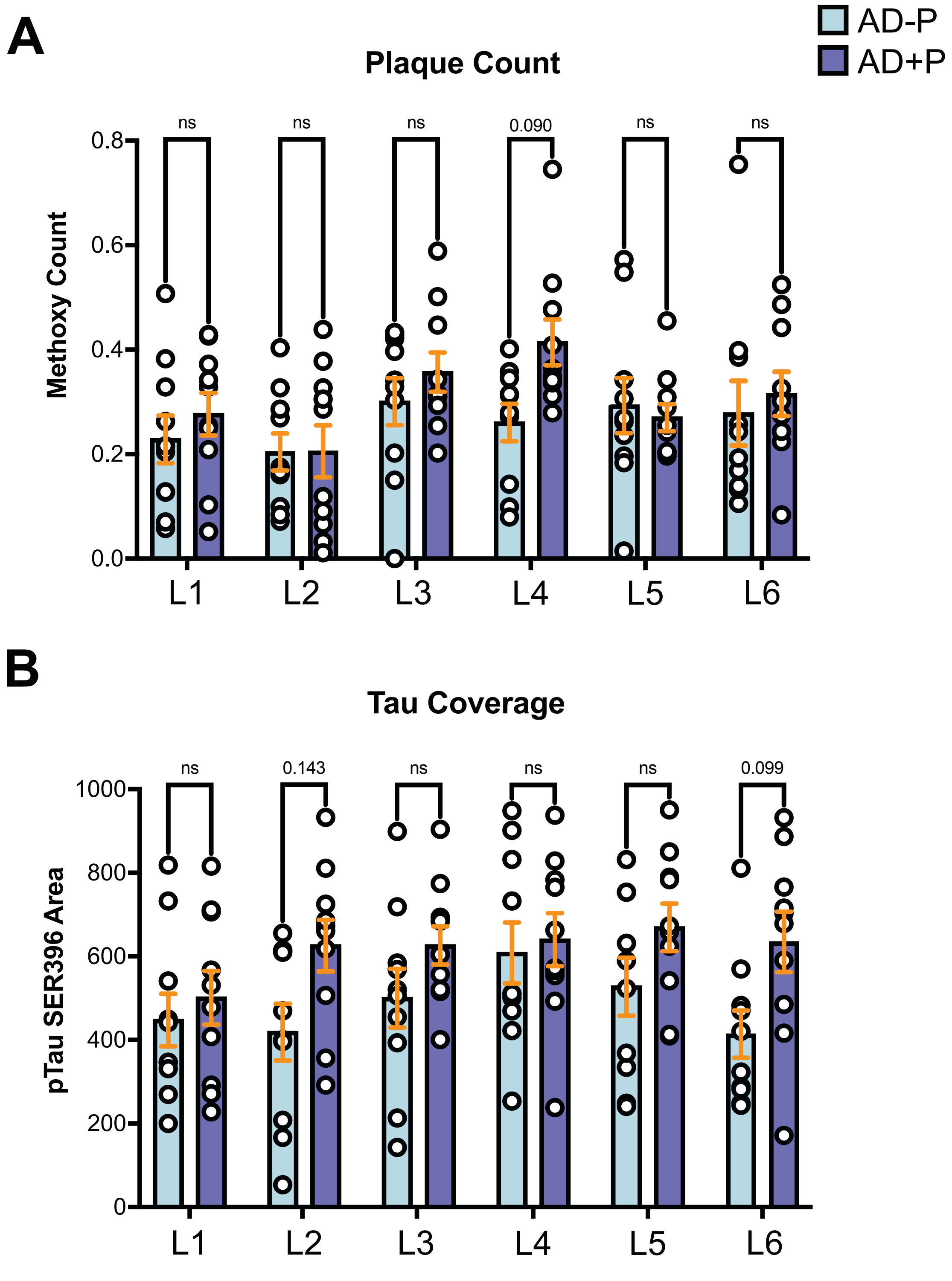


**Figure 1S: Histology for Neuropathological Changes in AD-P and AD+P.** An additional cohort of postmortem human brain tissue from 20 subjects (n= 10 AD-P and 10 AD+P). Error bar represents the SEM. Two-way ANOVA with Bonferroni's multiple comparison test. Values shown are normalized by area across layers and individuals captured in pixels using IMARIS.

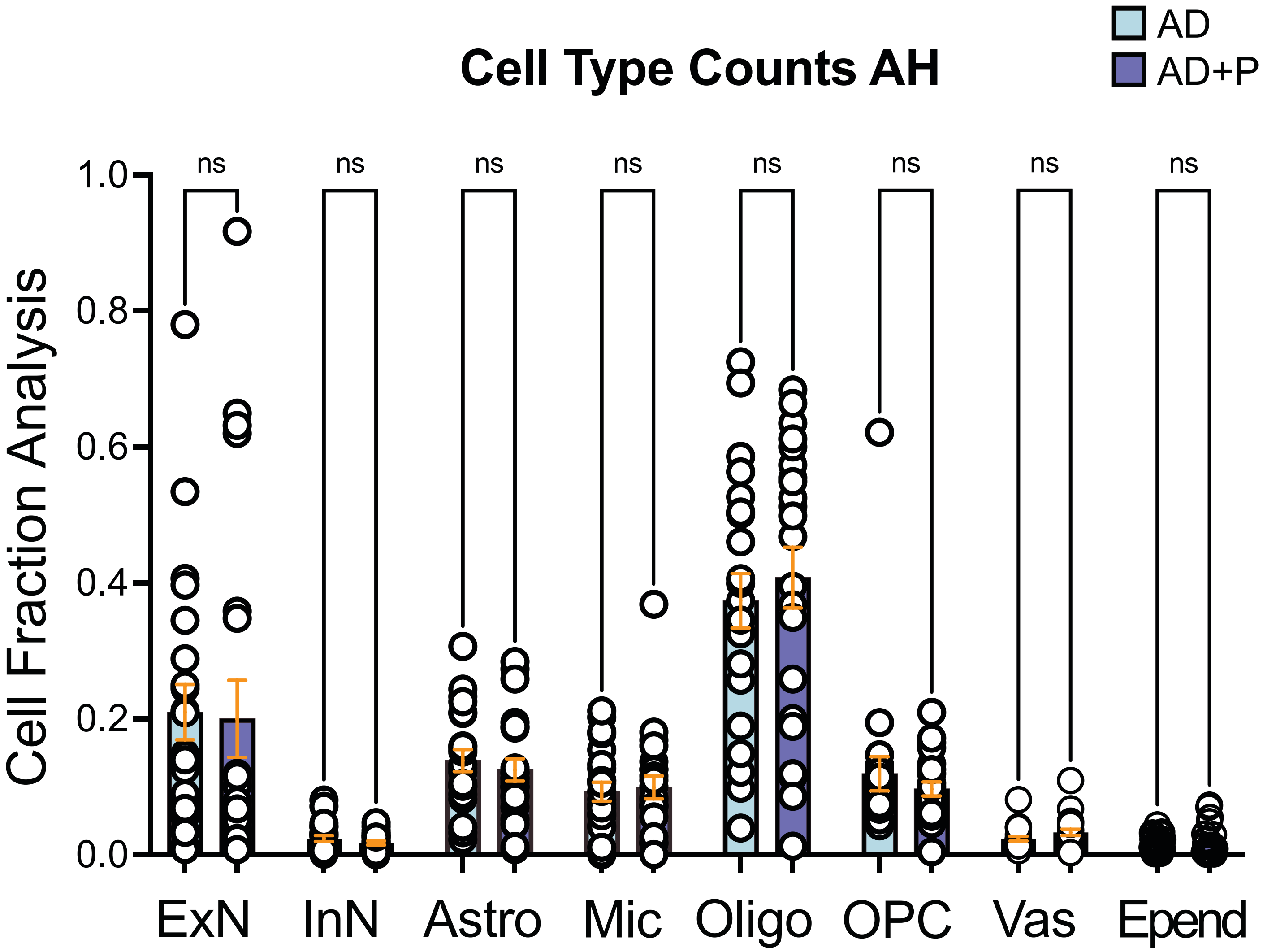


**Figure S2. Cell Fraction Analysis for Anterior Hippocampus in AD and AD+P.** Propeller analysis was used to determine statistical significance. Error bar represents the SEM. ns= not significant. n= 24 AD+P and 24 AD postmortem human brain tissue. AH: Anterior Hippocampus.

A

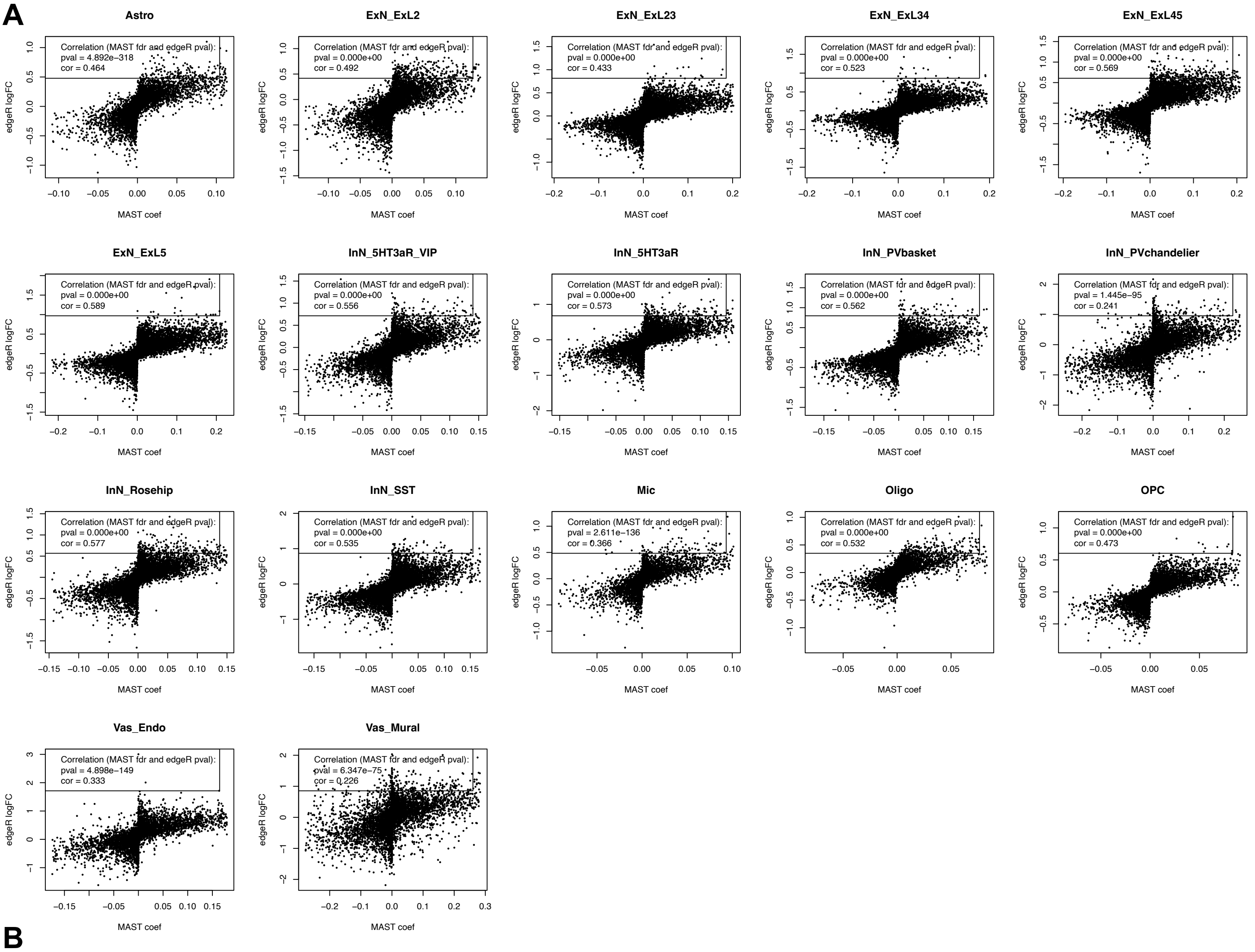


B

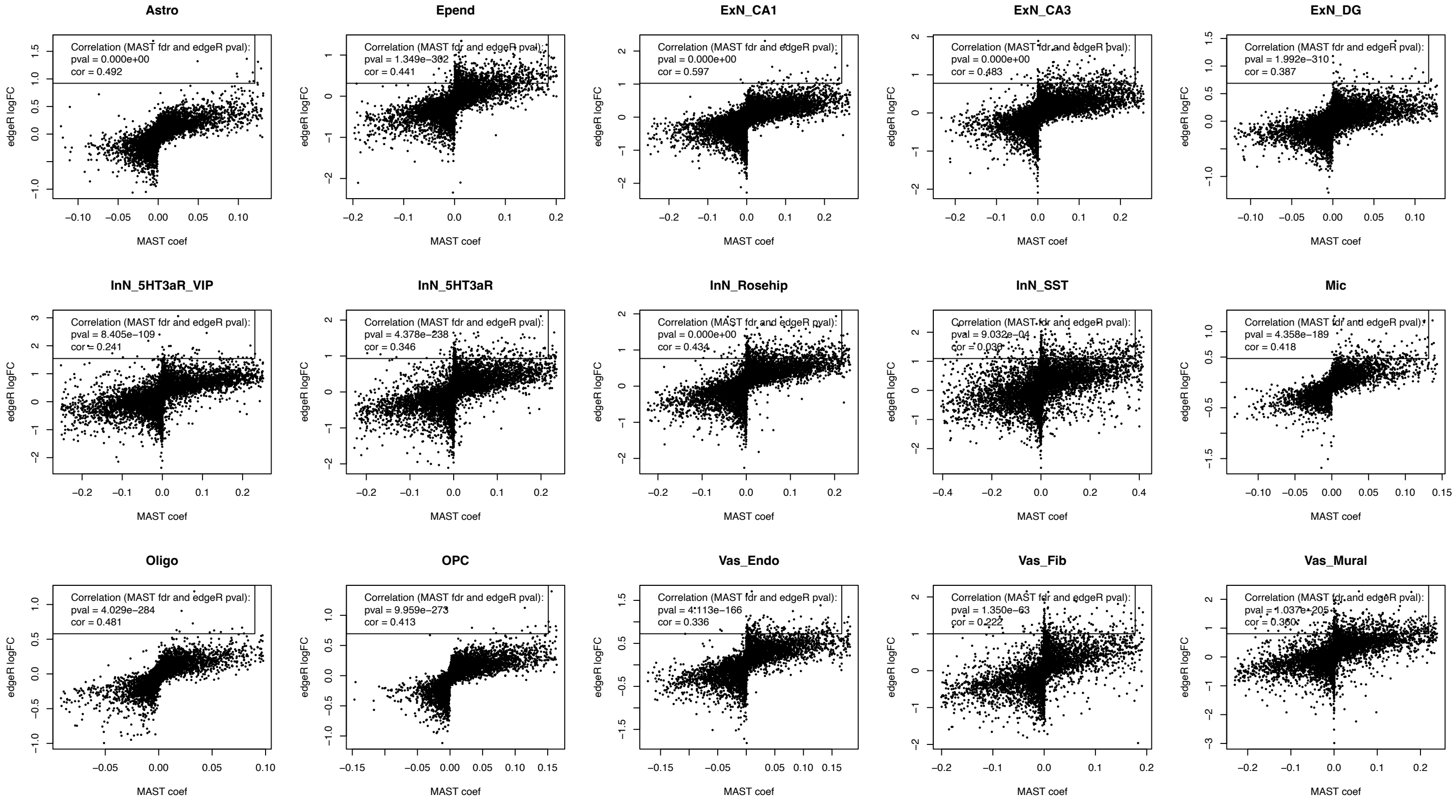


**Figure S3.Transcriptional Changes in AD with Psychosis.** A-B. Scatter plot to show the consistency of transcriptional changes identified by two methods in SF (A) and AH (B). The X-axis represents the coefficient calculated by MAST, while the Y-axis represents the log transformed fold change calculated by edgeR.

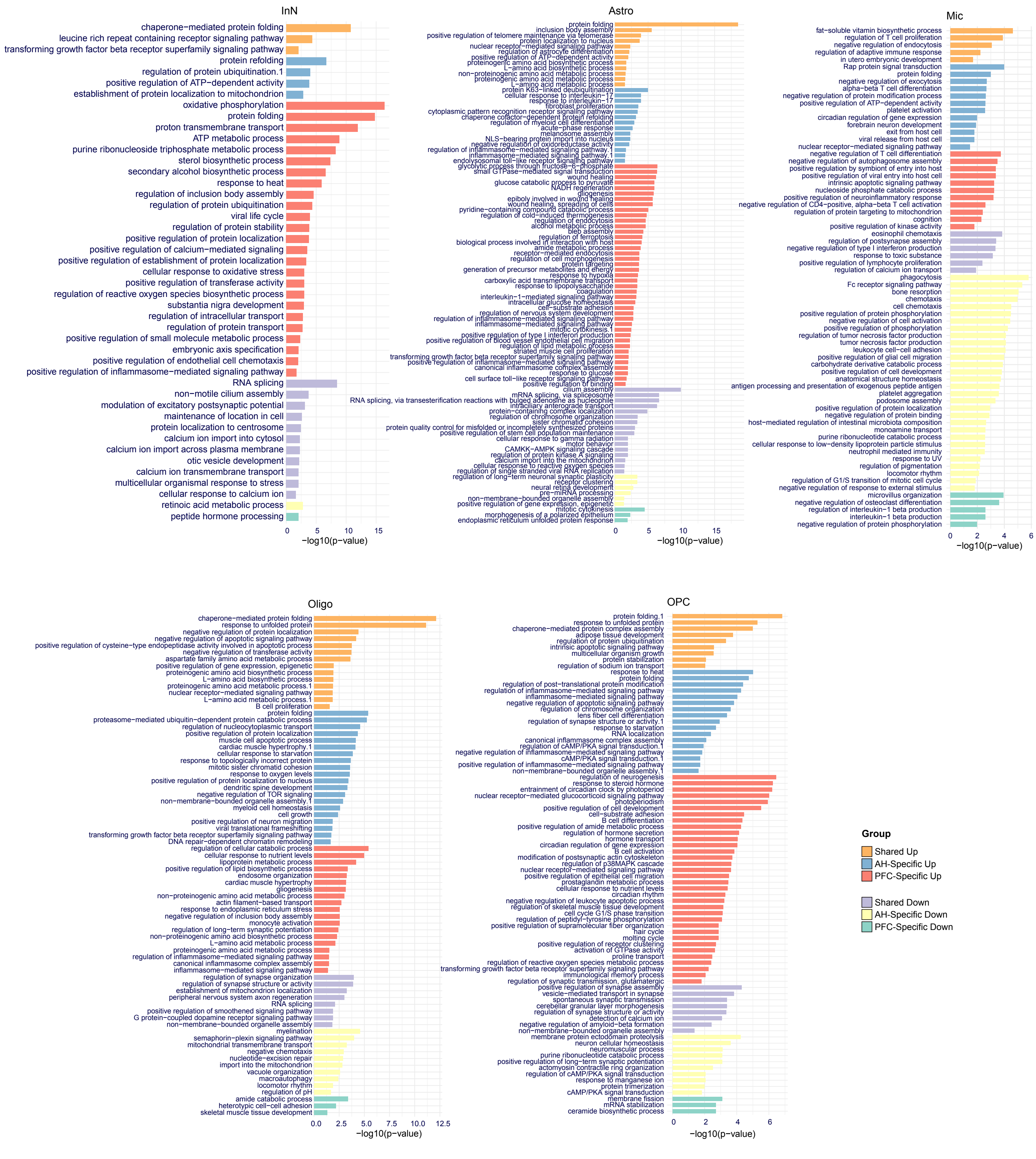


**Figure S4. Gene Ontology (GO) for Transcriptional changes in AD with psychosis.** Representative Gene Ontology from significantly enriched terms for five cell types.

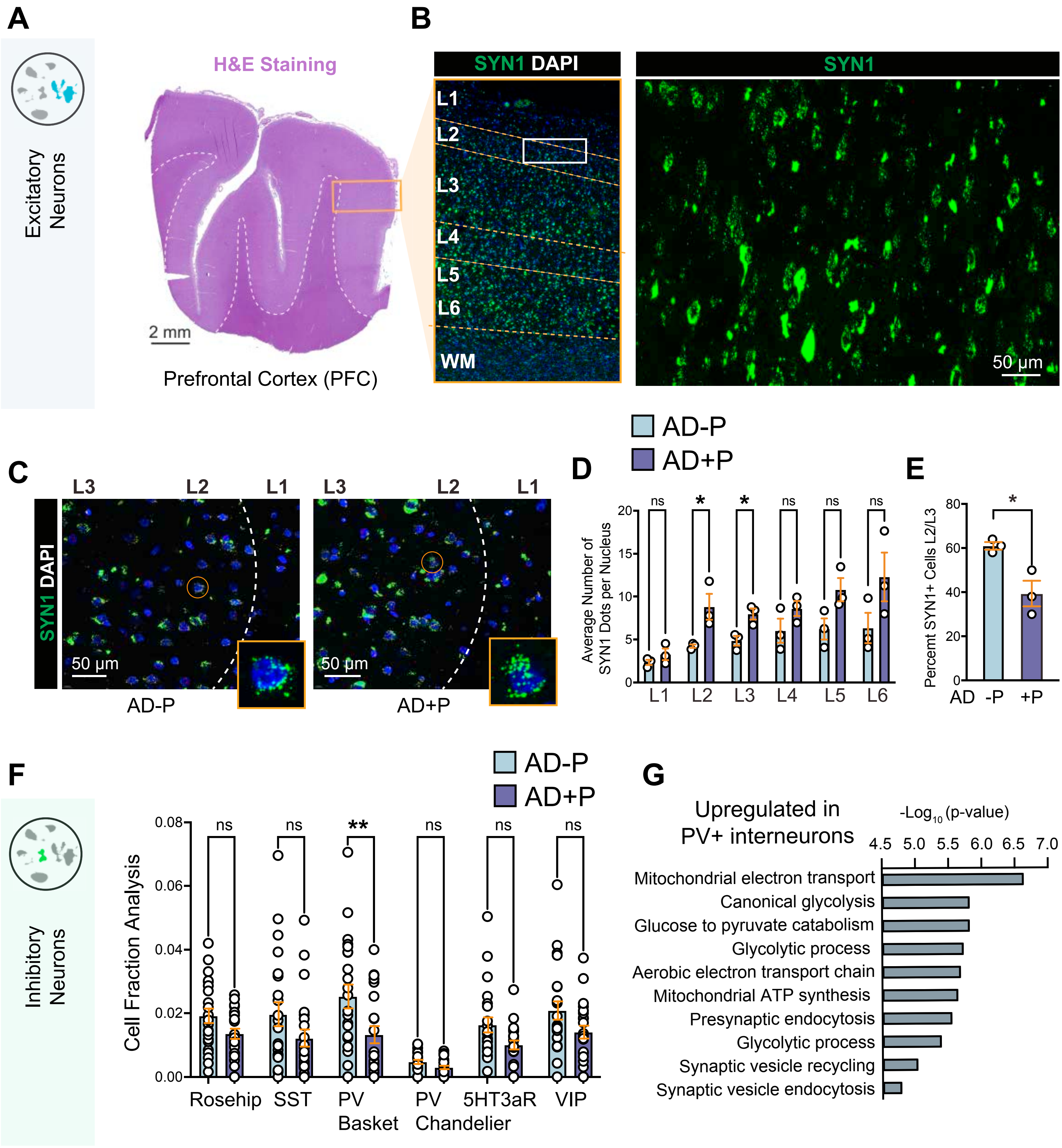


**Supplemental Figure S5. Compensatory mechanism evoked by surviving excitatory neurons and correlates of metabolic stress in PV+ interneurons.**

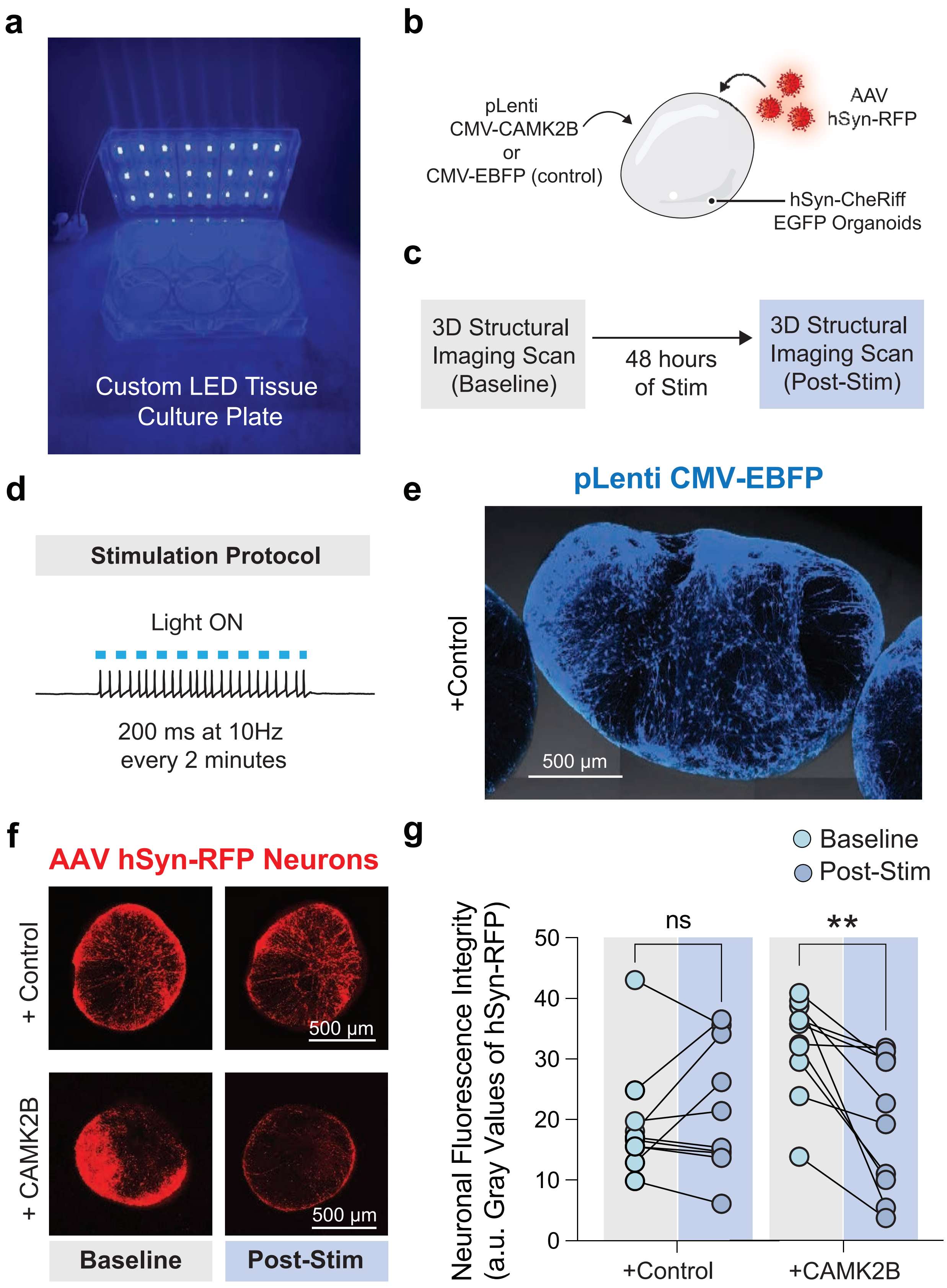


**Figure S6. CAMK2B Overexpression Renders Forebrain Organoids Vulnerable to Excitotoxicity.** A. Custom LED Tissue Culture Plate. B-C. Diagram of experimental design D. Stimulation protocol. E. Control organoids were tagged with pLenti CMV-EBFP. F. Organoids tagged with AAV hSYN-RFP before (baseline) and after stimulation (post-stim) and quantified (G). Paired t-test; **p-value<0.01, ns= not significant. n=10 control and 10 CAMK2B overpressing organoids.

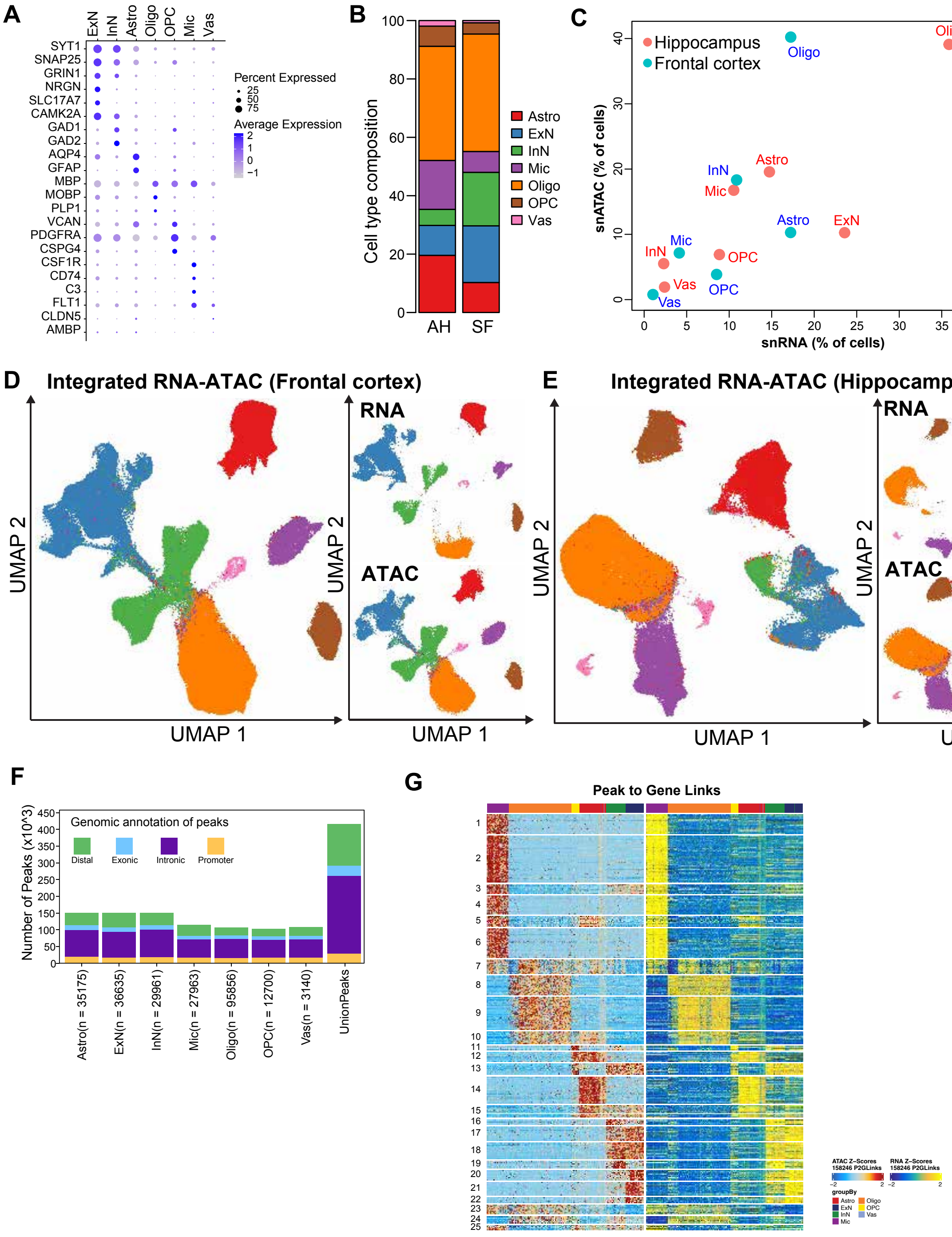


**Figure S7. Epigenetic Regulation of Cell-type-specific Gene Expression Changes. A.** Dotplot shows the gene activity score of well-known marker genes for major cell types in the human brain in snATAC-seq data. The color represents the activity level and the size of the dot represents the percentage of cells with activity. **B.** The composition of major cell types in snATAC-seq data in two brain regions, AH and SF. **C.** The comparison of cell fractions captured by snATAC-seq and snRNA-seq data in two brain regions. **D-E.** UMAPs to show the integrative analysis of snATAC-seq and snRNA-seq data in two brain regions. The UMAP on the left represents the joint map, and the right panel represents the nuclei from single modality. The nuclei were colored by major cell types.
**F.** The number of peaks per cell type and union set, colored by their genomic locations (distal, exonic, intronic or promoter regions). **G**. The heamap to show the links between distal peak to promoter peaks, representing the co-accessbility between distal enhancer and promoter. Each row represents one link and each column represents randomly selected nuclei (500 in total for visualization). The left panel represents the activity in s nATAC-seq, and the right plan represents the gene expression. Z-score values were used. Most of the links were cell-type specific.

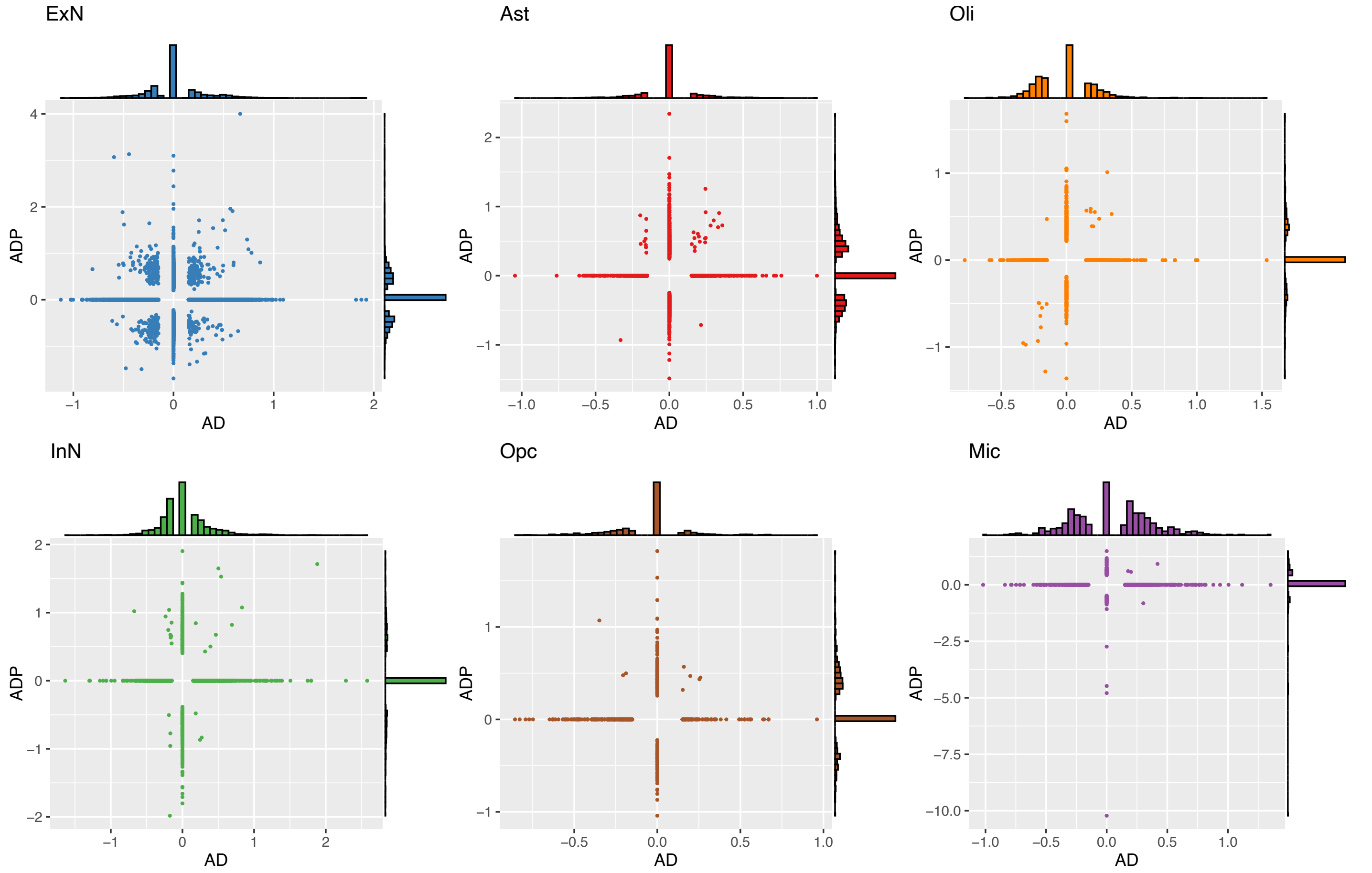


**Supplemental Figure S8. Limited concordance between canonical AD and AD+P transcriptional signatures across major brain cell types.** Scatter plots show the relationship between differential gene expression effect sizes in Alzheimer's disease (AD, x-axis), re-analyzed from Mathys et al. using the same differential expression framework applied in this study, and Alzheimer's disease with psychosis (AD+P, y-axis). Differential expression was computed relative to the appropriate control group for each dataset across all major brain cell classes, including excitatory neurons (ExN), inhibitory neurons (InN), astrocytes (Ast), oligodendrocytes (Oli), oligodendrocyte precursor cells (Opc), and microglia (Mic). Each point represents a gene tested for differential expression. Marginal histograms depict the distribution of effect sizes along each axis. This analysis illustrates both shared and divergent transcriptional responses between AD and AD+P across neuronal and glial populations.

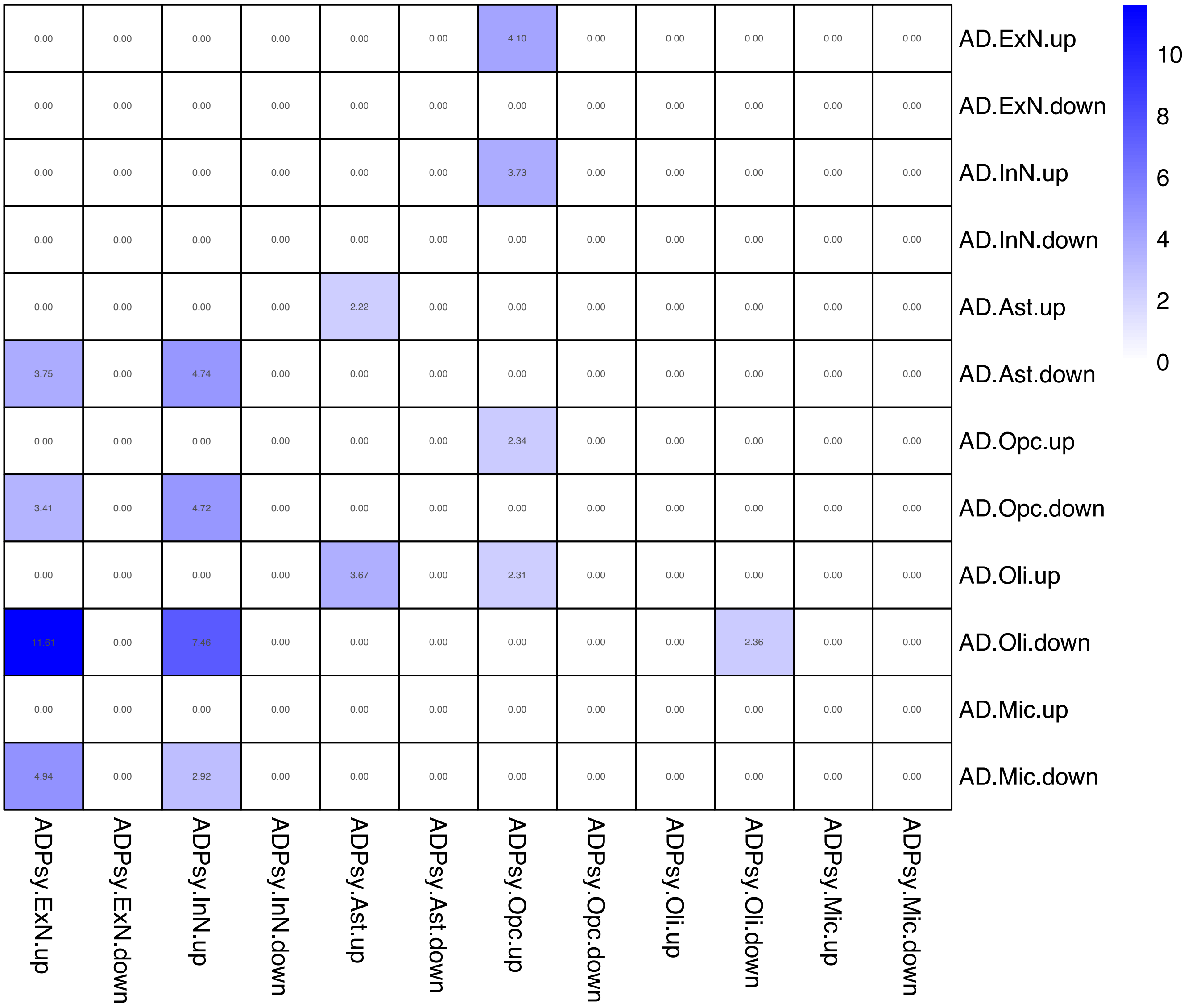


**Supplemental Figure S9. Cross-cell-type overlap of differential gene expression programs in AD and AD+P.** Heatmap depicts the degree of overlap between differentially expressed gene sets identified in Alzheimer's disease without psychosis (AD) and Alzheimer's disease with psychosis (AD+P) across major brain cell types, including excitatory neurons (ExN), inhibitory neurons (InN), astrocytes (Ast), oligodendrocyte precursor cells (Opc), oligodendrocytes (Oli), and microglia (Mic). Rows correspond to AD gene sets and columns to AD+P gene sets, each separated into up- and down-regulated categories. Cell values indicate the magnitude of gene overlap or enrichment between each gene set pair, as displayed by the color scale. This analysis highlights shared and distinct transcriptional programs across neuronal and glial populations in AD and AD+P.

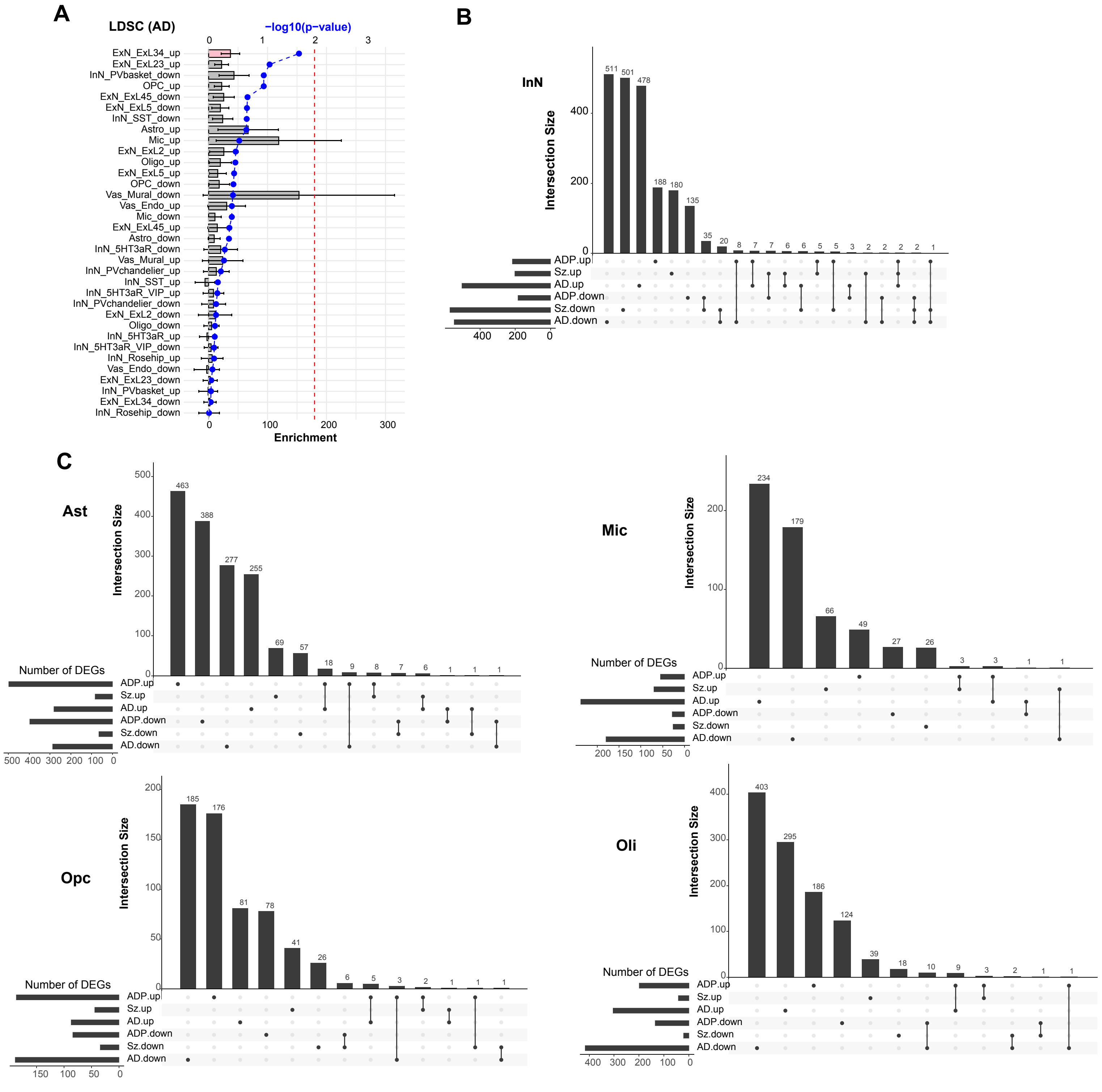


**Figure S10. Cell-type-specific Gene Expression Changes and their Association with Schizophrenia Genetics. A.** AD+P DEGs linked chromatin accessible peaks show the less enrichment of genetic variants of AD using LDSC analysis. The barplot shows the enrichment score, where pink ones represent the significance. The -log10 of p-value is shown by the blue point. The red dash line represents the stringent threshold with p-value < 0.01. **B-C**. The UpSet plot shows the overlap of DEGs in inhibitory neurons (**B**) and glial cell types (**C**) between Alzheimer's Disease (AD), Schizophrenia (Sz) and AD+P (ADP).
